# Alignment and Integration of Spatial Transcriptomics Data

**DOI:** 10.1101/2021.03.16.435604

**Authors:** Ron Zeira, Max Land, Benjamin J. Raphael

## Abstract

Spatial transcriptomics (*ST*) is a new technology that measures mRNA expression across thousands of spots on a tissue slice, while preserving information about the spatial location of spots. ST is typically applied to several replicates from adjacent slices of a tissue. However, existing methods to analyze ST data do not take full advantage of the similarity in both gene expression and spatial organization across these replicates. We introduce a new method *PASTE* (Probabilistic Alignment of ST Experiments) to align and integrate ST data across adjacent tissue slices leveraging both transcriptional similarity and spatial distances between spots. First, we formalize and solve the problem of pairwise alignment of ST data from adjacent tissue slices, or layers, using Fused Gromov-Wasserstein Optimal Transport (*FGW-OT*), which accounts for variability in the composition and spatial location of the spots on each layer. From these pairwise alignments, we construct a 3D representation of the tissue. Next, we introduce the problem of simultaneous alignment and integration of multiple ST layers into a single layer with a low rank gene expression matrix. We derive an algorithm to solve the problem by alternating between solving FGW-OT instances and solving a Non-negative Matrix Factorization (NMF) of a weighted expression matrix. We show on both simulated and real ST datasets that PASTE accurately aligns spots across adjacent layers and accurately estimates a consensus expression matrix from multiple ST layers. PASTE outperforms integration methods that rely solely on either transcriptional similarity or spatial similarity, demonstrating the advantages of combining both types of information.

**Code availability:** Software is available at https://github.com/raphael-group/paste

## 1 Introduction

Spatial transcriptomics (*ST*) is a new technology for measuring RNA expression in tissues while preserving spatial information [42]. ST involves placing a thin slice of tissue on an array covered by a grid of barcoded spots and sequencing the mRNAs of cells within the spots (Figure 1a). Early ST technologies [42] measured mRNA in up to 1000 spots, each spot containing 10 – 200 cells, while newer technologies such as the Visium technology from 10X Genomics [1] measure up to 5000 spots, each spot containing up to 10 cells. ST has been used to study cancer tissue (e.g. breast [42], prostate [5], melanoma [45], pancreas [34], carcinoma [25]), diseased tissues (e.g. Alzheimer’s [10] and gingivitis [31]) and healthy tissues (e.g. mouse olfactory bulb [42], human heart [3], spinal cord [33]), as well as other applications. Multiple computational methods have been introduced to analyze ST data, including the identification of spatial patterns of gene expression [42, 30], spatially distributed differentially expressed genes [44, 5, 2] and spatial cell-cell communication patterns [2, 9].

**Figure 1:**
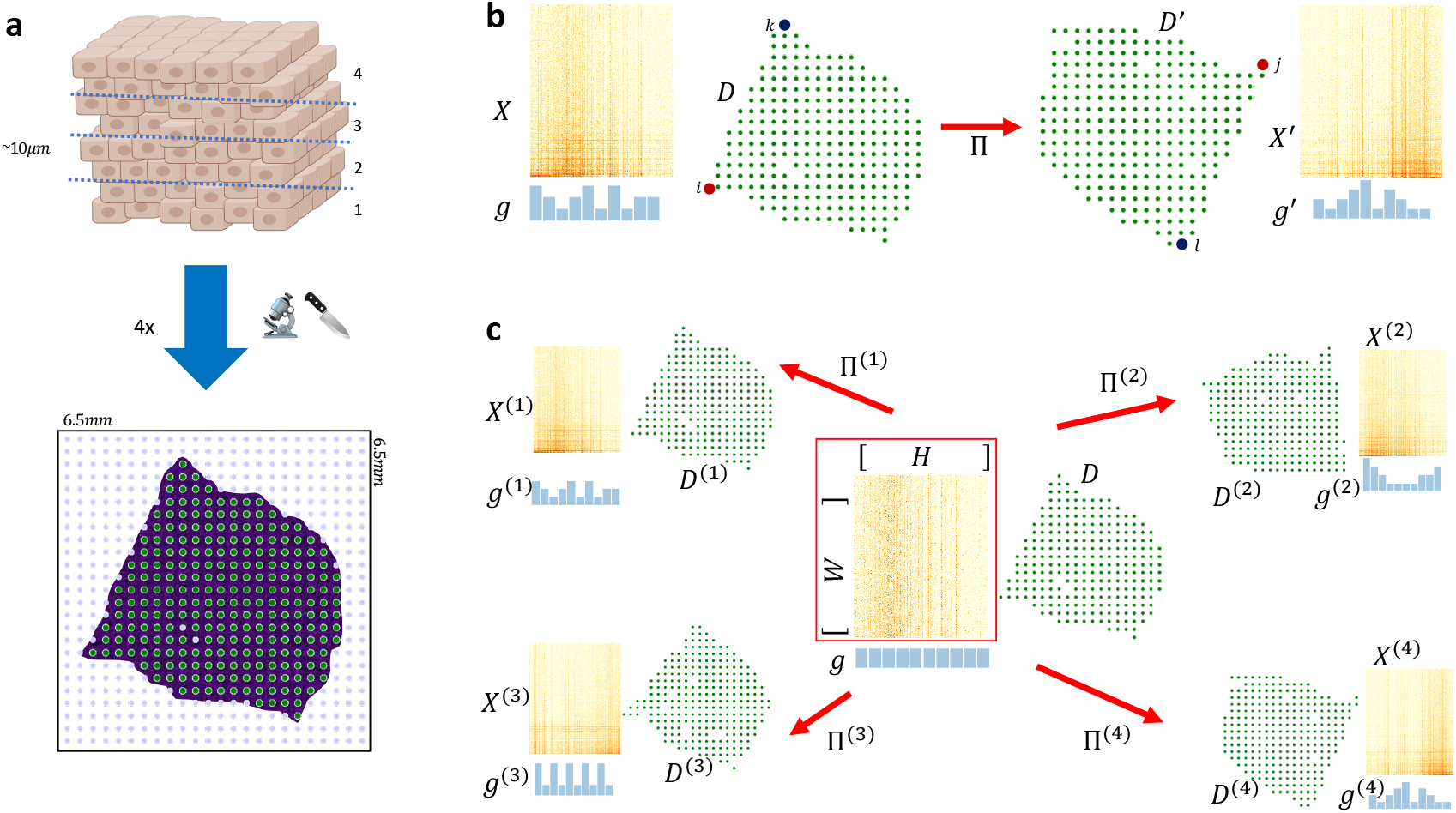
(a) A spatial transcriptomics (ST) experiment starts by dissecting a tissue into narrow (10 – 20*μm*) layers. Each layer is placed on a 2D grid of barcoded spots, and mRNA expression of each spot is measured along with the spatial coordinates of each spot. Only a fraction of spots (green) contain tissue cells, with other spots (blue) not covered by a tissue. (b) The Pairwise Layer Alignment Problem aims to find a mapping Π = [*π_ij_*] between spots in one layer and spots in another layer while preserving the gene expression and the spatial distances of mapped spots. (c) The Center Layer Integration Problem aims to infer a“center” layer consisting of a low rank expression matrix *X* = *WH* and a collection Π^(1)^,… Π^(*t*)^ of mappings from the spots of the center layer to the spots of each input layer.

While many ST studies generate data from multiple adjacent tissue layers, nearly all current ST analysis techniques either analyze only individual layers [34, 15] or pool only gene expression data across layers without considering the spatial coordinates [5, 25]. However, by ignoring shared spatial information across layers, such methods lose power in their analyses. A recently developed software package named STUtility [4] aligned stacked ST experiments using their histological images by identifying transformations of the images that match the tissue edges. However, STUtility does not consider the gene expression data or locations of spots. Thus, depending on the topology of the tissue, STUtility may fail to automatically align the images and the user must resort to manual image alignment. More importantly though, STUtility does not output a mapping between spots that can be used for downstream analysis.

Multiple methods have been introduced to integrate data from single-cell assays such as scRNA-seq, ATAC-seq, etc. [22, 43, 23, 32, 13, 48], that do not include spatial information. While these methods could be applied to ST data by ignoring the spatial coordinates of the spots, the spatial coordinates provide a rigid structure to the ST data and cannot simply be treated as additional features. Moreover, due to differences in the dissection of the tissue layers and their placement on the array, the absolute values of the spatial coordinates cannot be easily compared across layers. Therefore, integration of ST that preserve both gene expression and spatial data is nontrivial.

We introduce *PASTE* (Probabilistic Alignment of ST Experiments), a method to align and integrate multiple tissue layers from an ST experiment using information from both gene expression and spatial coordinates. First, we formalize the problem of probabilistic pairwise alignment of adjacent layers based on transcriptional and spatial similarity using Fused Gromov-Wasserstein Optimal Transport (*FGW-OT*)[46]. This enables the reconstruction of a 3D spatial dataset by sequentially aligning multiple adjacent ST layers. However, since the thickness of each layer is relatively small in comparison to the size of spots and the spacing between spots, this 3D structure is limited. Therefore, we formalize the problem of finding a “center” layer that integrates multiple ST layers by combining FGW-OT Barycenter formulation [46] with an assumption that the gene expression matrix is low rank. We prove that the restricted optimization problems reduce to either an FGW-OT problem or a Non-negative Matrix Factorization (NMF) [28] problem, and give a block coordinate descent algorithm for finding a center layer. The combined inference of a center layer has the potential to increase the power of downstream analysis relative to many of the current ST analysis methods that analyze single layers individually.

We demonstrate the utility of PASTE on both simulated ST datasets and a recently published squamous cell carcinoma dataset [25]. We show on simulated data that using both gene expression data and spatial coordinates when aligning pairs of adjacent layers or combining multiple layers is superior to using either data type alone. We show that PASTE provides better accuracy in aligning spots across layers and in recovering the gene expression patterns of the tissue. Finally, we show on the squamous cell carcinoma ST dataset that PASTE aligns layers while preserving the spatial relationships between different cell type compositions, thus providing additional validation. Furthermore, we show that combining multiple ST layers improves downstream analysis by enabling more spatially coherent clustering than either ST layer alone.

## 2 Methods

We start by describing how we represent ST layers and define some required notations in Section 2.1. We pose the problems of aligning a pair of ST layers or combining multiple layers into a single layer in Sections 2.2 and 2.3.

### 2.1 Representing spatial transcriptomics layers

Let *M* = [*m_ij_*] be a matrix, let *m_i_*. be a vector corresponding to the *i*^th^ row of *M*, and let *m._j_* be a vector corresponding to the *j*^th^ column of *M*. We use 1_*n*_ to denote a column vector of length *n* containing all ones. For a vector *v* of length *n*, we denote by diag(*v*) an *n* × *n* matrix with diag(*v*)_*ii*_ = *v_i_* and zero for all other entries and denote by 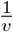 an a vector with the element-wise inverse of the values of *v*. We denote by Tr(*M*) = ∑*m_ii_* the trace of a square matrix *M*.

The result of an ST *experiment* is a pair (*X, Z*) of matrices, where 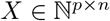 is a *p* genes by *n* spots expression matrix and 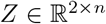 is the coordinate matrix of the spots. That is, 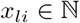 is the transcript count for gene *l* in spot *i* and *z._i_* is the coordinate vector of spot *i* on the array. Since the placement and orientation of the tissue on the array are arbitrary, we find it more convenient to represent only the relative location of the spots. Therefore, instead of the actual spots locations *Z*, we use the spot distance matrix 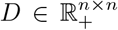, where *d_ij_* = ∥*z._i_* − *z._j_*∥ is the spatial distance between spot *i* and *j*. While this transformation is not reversible, it has an advantage of being invariant to the translation or rotation of the tissue on the array.

In addition to the gene expression and distance matrices, we assume that each tissue spot *i* has a weight *g_i_* > 0 representing its relative importance compared to the other spots^1^. These weights encode prior information on the spots such as the relative number of cells in the spot, the presence of a cell surface marker in the spot or an importance score of the spot based on pathological examination of the tissue. We assume these weights are normalized so ∑_*i*_ *g_i_* = 1 and thus *g* = (*g*_1_,…, *g_n_*) is a *distribution* over the spots. If no prior information is given on the spots, we use a uniform distribution 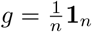 over the spots.

A spatial transcriptomics *layer* of *n* spots over *p* genes is described by a triplet (*X, D, g*) where 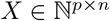 is a gene by spot expression matrix, 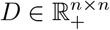 is the spot pairwise distance matrix and *g* is a distribution over spots. We call the column vector *x._i_* the *expression profile* of spot *i*.

### 2.2 Pairwise Alignment of ST layers

We start by defining the problem of mapping/aligning a pair of ST layers. Our goal in this problem is to find a mapping between spots in the two layers such that spots mapped to one another have similar expression profiles and the spatial relationship of spots within each layer is preserved by the mapping. Ideally one would want a one-to-one mapping (or perfect matching) between spots in the different layers. However, such a matching of the spots is not always feasible and may not even be a suitable due to the intrinsic variations of the ST experiment. First, the number of spots and their locations in the tissue in each layer may be different due to differences in dissections of the tissue, the placement on the array and the sequencing coverage of spots. Therefore, there may be spots in one layer that have no direct match to a spot in the other layer. Second, since the placement of the tissue with respect to the fixed position of the spots on the array changes between layers, the true position in the tissue of some spots in one layer may actually fall in between several spots in the other layer.

We propose to formalize the problem of aligning spots across layers as a many-to-many fractional mapping problem where the weight of each spot in one layer is allowed to be split to several spots. Inferring such a fractional mapping is the subject of the field of Optimal Transport (OT) Theory [49]. Recent advancements in optimal transport theory enable efficient algorithms [11, 12, 38] and flexible formulations [39, 46], thus facilitating its use in numerous applications. In particular, OT has been applied in single cell analysis to infer developmental trajectories [40], reconstruct spatial expression [36], infer cell-cell communication [9] and integrate multi-omic datasets [13].

Let (*X, D, g*) and (*X′, D′, g′*) be two layers of *n* and *n′* spots respectively over the same *p* genes. We say that a matrix 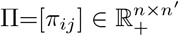 is a *mapping* between the two layers if for all spots *i* in the first layer ∑_*j*_ *π_ij_* = *g_i_* and for all spots *j* in the second layer ∑_*i*_ *π_ij_* = *g′_j_*. We denote by Γ(*g, g′*) the set of all mappings between the two layers. We say that a spot *i* in one layer and spot *j* in the other layer are *mapped/aligned* if *π_ij_* > 0.

We define the Pairwise Layer Alignment Problem as finding a mapping between spots in one layer and spots in another layer that takes into account both the inter-layer transcriptional dissimilarity and the intra-layer spatial distance structure (Figure 1b). We formalize this problem as a Fused Gromov-Wasserstein Optimal Transport problem [46] with an *expression cost function c* : 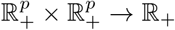 that measures a non negative cost between the expression profiles of two spots over all genes.

#### Pairwise Layer Alignment Problem.

*Given layers* (*X, D, g*) *and* (*X′, D′, g′*) *containing n and n′ spots respectively over the same p genes, an expression cost function c and a parameter* 0 ≤ *α* ≤ 1, *find a mapping* Π ∈ Γ(*g, g′*) *minimizing the following transport cost:*

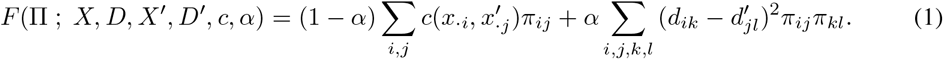

The parameter *α* balances between the transcriptional dissimilarity between spots induced by the mapping and the spatial distance similarity of the spots induced by the mapping. When *α* = 0 only the gene expression data is taken into account, while when *α* =1 only the spatial coordinates are taken into account. Note that the problem is invariant to translation or rotation of the coordinates of any layer since the transport cost depends only on distances *D* and *D′* within a layer and not on the absolute spatial coordinates. In addition, note that if *α* = 0, *n* = *n′* and 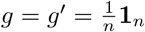, the problem reduces to finding a minimum weight bipartite perfect matching of the spots across layers [6].

We solve the Pairwise Layer Alignment Problem using the iterative conditional gradient algorithm described in [46] for the fused Gromov-Wasserstein optimal transport problem. This algorithm takes *O*(*n*^2^*n′* + *nn′*^2^) operations per iteration.

Solving the Pairwise Layer Alignment Problem between a adjacent layers is useful for reconstructing a 3D spatial representation of tissue. Namely, given a series (*X*^(1)^, *D*^(1)^, *g*^(1)^),…, (*X*^(*t*)^, *D*^(*t*)^, *g*^(*t*)^) of sequential layers we find the mapping Π^(*k*)^ between adjacent layers *k* and *k* + 1 for every *k* =1,…, *t* − 1. To project all layers to the same spatial coordinate system we use the mappings to solve a generalized weighted Procrustes problem [50, 27]. That is, we seek to project the spatial coordinates *Z*^(*k*+1)^ of layer *k* + 1 to the spatial coordinates *Z*^(*k*)^ of layer *k* by finding a translation vector 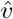 and rotation matrix 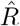 that minimize the weighted distances between mapped spots (Supplementary Section S1.2). Formally, we solve

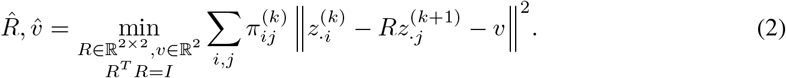

The projected spatial coordinates of spot *j* in layer *k* + 1 are then 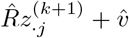. To solve the weighted Procrustes problem given by Equation 2 we use SVD (Supplementary Section S1.2).

### 2.3 Integration of multiple ST layers

A natural generalization of pairwise alignment of two layers is the integration of multiple layers into a single consensus layer. This integration leverages transcriptional and spatial similarities simultaneously across all layers, and thus can overcome variability in individual layers due to varying sequencing coverage, tissue dissection, or tissue placement on the array. A potential disadvantage is that a single consensus layer does not yield a 3D reconstruction of the tissue. However, in current ST datasets the thickness of each tissue slice (10-20 microns) is smaller than the diameter of spots (100 microns in ST and 55 microns in Visium) and the spacing between spots (100-200 microns). Thus, the additional information obtained from a 3D reconstruction is relatively limited, and in practice the advantages of multi-layer integration may outweigh the disadvantage of not obtaining a 3D reconstruction.

We define the problem of integrating multiple ST layers (e.g. from replicate experiments or adjacent tissue slices) into a single center layer that is similar to the individual layers in both gene expression and spatial relationships between spots (Figure 1c). This problem is analogous to the “star alignment” problem in multiple sequence alignment [21], but with added complexity resulting from the two dissimilarity measures. The combined inference of a center layer has the potential to increase the power of downstream analysis of ST relative to many of the current ST analysis methods that analyze single layers individually. Furthermore, summarizing the ST layers into one consensus layer can pool information across layers and overcome errors in individual layers.

We define the integration problem under a few reasonable biological and computational assumptions. First, we assume that the given ST layers are very similar to each other and thus can be summarized with a single layer. This assumption is again motivated by the fact that the thickness of an ST layer is small relative to the diameter of a spot and the spacing between spots. Therefore, ST layers from the same tissue are often referred to as technical replicates [5, 34, 25]. Second, we assume that spatial coordinates and the distribution over spots in our center layer are known in advance, up to rotation or translation. This is a reasonable assumption since the spatial coordinates on the array are fixed by the technology. Finally, we assume that the expression matrix of the center layer is low rank. This is a widely used assumption in both in single cell RNA-seq analysis and ST analysis and corresponds to the biological assumption that cells/spots often occupy a limited number of cell types or cell states [29, 35, 24]. In addition, in most ST experiments, the number of ST layers per tissue is small (2-4) and the gene expression matrices are relatively sparse (≥75% zeros), and thus estimation of a full rank gene expression matrix would be prone to overfitting.

We formalize the problem of finding a center ST layer by combining the ideas of fused Gromov-Wasserstein barycenter [46] and Non-Negative Matrix Factorization [28]. NMF has been shown to be useful in single-cell RNAseq analysis both as a method to impute missing values (“dropouts”) and as a dimensionality reduction technique [41, 53, 17]. In the Center Layer Integration Problem similar to the fused Gromov-Wasserstein barycenter problem – we seek to find a center ST layer that minimizes the weighted sum of distances to a given set of input ST layers, where the distance between layers is calculate by the minimum value of the Pairwise Layer Alignment Problem objective across all mappings. However, unlike the fused Gromov-Wasserstein barycenter problem, we also require the consensus gene expression matrix to be non-negative and low rank (Figure 1c).

#### Center Layer Integration Problem.

*Given layers* (*X*^(1)^, *D*^(1)^, *g*^(1)^),…, (*X*^(*t*)^, *D*^(*t*)^, *g*^(*t*)^) *containing n*_1_,…, *n_t_ spots, respectively over the same p genes, a spot distance matrix* 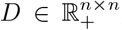, *a distribution g over n spots, an expression cost function c, a distribution* 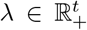, *and parameters* 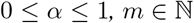 *and, find an expression matrix X* = *WH where* 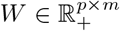 *and* 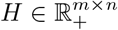, *and mappings* Π^(*q*)^ ∈ Γ(*g, g*^(*q*)^) *for each layer q* = 1,…, *t that minimize the following objective:*

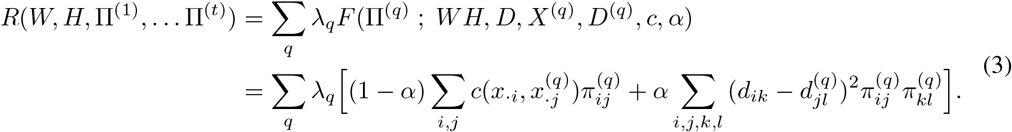

We solve the Center Layer Integration Problem, we propose a Block Coordinate Descent algorithm (Algorithm 1). This algorithm alternates between optimizing the mappings Π^(1)^,…,Π^(*t*)^ given the current values of *W, H* and optimizing *W, H* given the current mappings Π^(1)^,… Π^(*t*)^. The problem of finding the optimal mappings Π^(1)^,… Π^(*t*)^ given *W* and *H* reduces to solving the Pairwise Layer Alignment Problem between the center layer (*WH, D, g*) and each layer (*X*^(*q*)^, *D*^(*q*)^, *g*^(*q*)^) separately. Similarly, the problem of finding the optimal *W* and *H* given the current mappings Π^(1)^,…, Π^(*t*)^ reduces to a new problem we call the Center Mapping NMF Problem. We show below that the Center Mapping NMF Problem can be interpreted as a maximum likelihood optimization and prove it is equivalent to a weighted NMF problem.

#### Algorithm 1

BCD for Center Layer Integration Problem

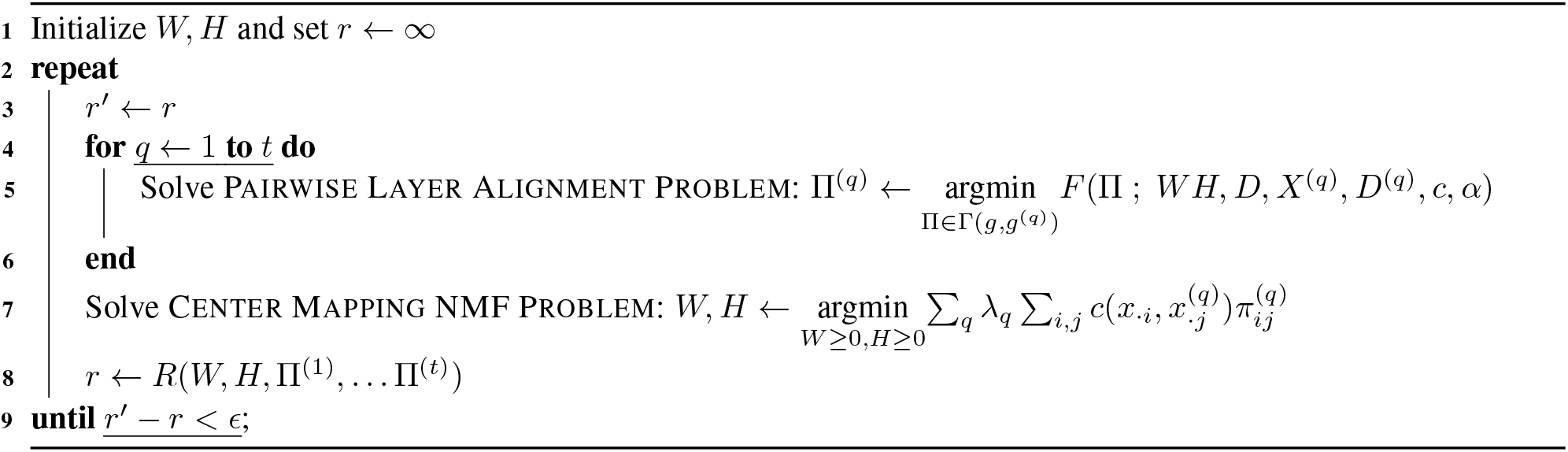

#### Center Mapping NMF Problem.

*Given t expression matrices* 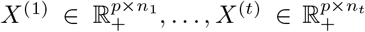, *t mapping matrices* Π^1^ ∈ Γ(*g,g*^(1)^),… Π^(*t*)^ ∈ Γ(*g, g*^(*t*)^), *an expression cost function c, a distribution* 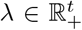 *and parameters* 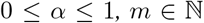 *find two low rank matrices* 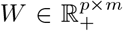 *and* 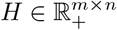 *such that X* = *WH minimizing the following objective:*

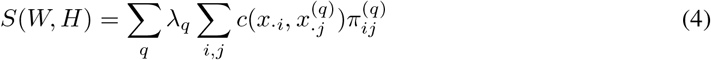

We analyze the Center Mapping NMF Problem for two commonly used expression cost functions [28]: (1) the Euclidean distance *c*(*u, v*) = ∥*u − v*∥^2^ and (2) the KL divergence 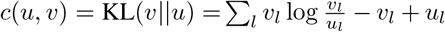. While the KL divergence is not symmetric and therefore not a distance measure, it has the advantage of having a probabilistic interpretation as the likelihood of a Poisson count model [20]. Thus, it has been used in the analysis of count data matrices such as sc-RNAseq [14, 47, 16]. Hence, solving the Center Mapping NMF Problem is motivated by finding a low rank expression matrix *X* = *WH* that maximizes the likelihood of the following generative model when 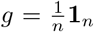 and 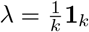:

- The random variables of the number of transcripts 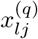 of a gene l in spot *j* in layer *q* are independent given *X*, Π^(1)^,…, Π^(*t*)^.
- The number of transcripts 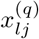 of a gene *l* in spot *j* in layer *q* given that it was generated from spot *i* in the consensus layer has a distribution 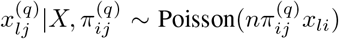. Therefore, the total number 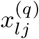 of transcripts of a gene *l* in spot *j* in layer *q* is 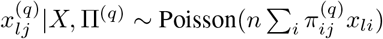.

The negative log likelihood of the model is

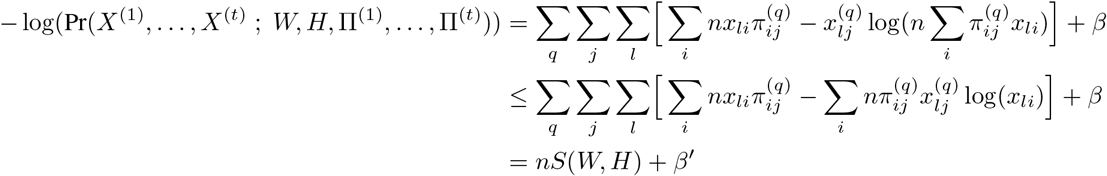

where *β* and *β′* are constants independent of *W* and *H*. The second transition follows from Jensen’s inequality. Therefore, minimizing *S*(*W,H*) with respect to *W, H* maximizes the likelihood of this probabilistic model.

The Center Mapping NMF Problem is equivalent to the problem of finding a weighted NMF [7] of the matrix 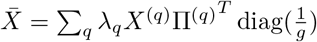 as stated in the following theorem:

#### Theorem 1.

*Let* 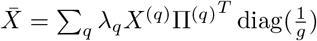 *and X* = *WH*. We have,

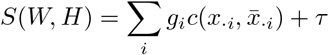

*where c*(*u, v*) = ∥*u − v*∥^2^ *or* 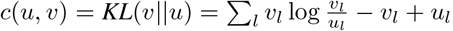, *and τ is a constant that does not depend on W, H*.

The proof of Theorem 1 is given in supplementary Section S1.

As a result of Theorem 1 we can solve the Center Mapping NMF Problem using an algorithm for weighted NMF such as the iterative update scheme of [7]. When the distribution *g* of the spots in the center layer is a uniform distribution, i.e. 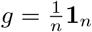, the Center Mapping NMF Problem reduces to a traditional NMF [28] of the matrix 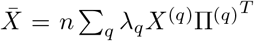. While in our problem formulations we assumed the low rank matrices *W, H* are non-negative, Theorem 1 does not use this assumption. Therefore, we can use other factorization techniques such as PCA or generalized PCA [47] in our problems and algorithms.

### 2.4 Implementation details and parameter selection

We implemented the algorithms described in Sections 2.2 and 2.3 in a software package called PASTE. To solve the Pairwise Layer Alignment Problem, we use the fused Gromov-Wasserstein optimal transport algorithm implementation from the Python Optimal Transport library [19]. To solve the Center Layer Integration Problem, we implemented Algorithm 1 in Python. We initialize *W, H* in Algorithm 1 by running NMF on one of the input layers and stop the iterations when the improvement in the objective function is less than *ϵ* = 10^−3^. In all analyses described below, we use a uniform distribution 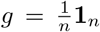 for the spots in a layer with *n* spots, give all layers an equal weight 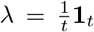 in the Center Layer Integration Problem and use *m* = 15 dimensions for NMF. We use the KL divergence to calculate the expression cost between spots. For both problems, we use *α* = 0.1 (unless otherwise specified) based on our performance on simulated data (Supplementary Figure S2). Before running the algorithms, layers are preprocessed such that genes with non-zero expression in fewer than 15 spots or genes that are not expressed in all input layers are removed.

Pairwise layer alignment using PASTE takes ≈9-12 seconds on layers with 260-270 spots and 8000 genes. The center layer integration takes ≈90 seconds to integrate three layers with the same numbers of spots and genes. PASTE was run on an Alienware Aurora R9 with an Intel^®^Core™ i9-9900 processor.

## 3 Results

We evaluate PASTE on both simulated ST data and ST data from squamous cell carcinoma (SCC) [25]. In Section 3.1, we describe our ST simulation and show that using both transcriptional and spatial data enables PASTE to correctly align and integrate ST layers. In Section 3.2, we apply PASTE to SCC data and show that it is able to map spots across layers while preserving spot cluster annotations.

### 3.1 Analysis on Simulated Data

We first evaluated PASTE on simulated ST data where the correspondence between spots across layers is known. To ensure that the both expression and topology of the simulated data are biologically reasonable we used real samples of ST from four layers of a breast tumor [42] to generate our simulated ST data. Each layer in this dataset consists of 251-264 spots and 7453-7998 genes (Supplementary Figures S1 and S9).

We simulate a new ST experiment (*X′, Z′*) with *n′* tissue spots from a given ST experiment (*X, Z*) with *n* tissue spots by perturbing both the gene expression and spatial data as follows. First, we assume that the locations 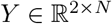 of all spots on the array are known and that tissue spots will only be generated from these locations. This is a reasonable assumption since the spot locations on the array are fixed. For each spot *i* in the original ST experiment we generate new transcript counts and spatial coordinates using the following procedure which is governed by a *coverage variability factor η* controlling the variance of the number of read counts per spot and a parameter *θ* controlling the spatial rotation of the tissue on the array.

1. Select *k_i_* total read counts according to *k_i_* ~ NegativeBinomial(*μ, νη*), where 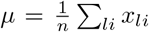 is the empirical mean spot total read count, 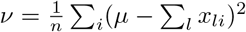 is the empirical variance of the spots total read count.
2. Generate an expression profile 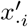 for spot *i* according to 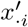 ~ DirichletMultinomial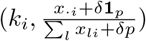, where *δ* = 1 is a small pseudo-count.
3. Generate rotated coordinates *v._i_* = *z._i_*Θ, where Θ is a rotation matrix with an angle *θ*. Then, coordinates 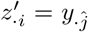 are mapped to the closest spot on the array grid 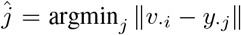. If the grid spot 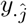 was already mapped to a previous tissue spot, spot *i* is discarded.

Note that step 3 may result in a simulated layer with fewer spots than the original layer. For instance, the original layer 1 (Figure 2a) contains a total of 254 spots while a simulated layer with 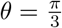 (Figure 2b) contains only 220 spots. We denote by *i* ~ *j* a spot *i* in the original layer and a spot *j* in the simulated layer that are mapped to one another.

**Figure 2:**
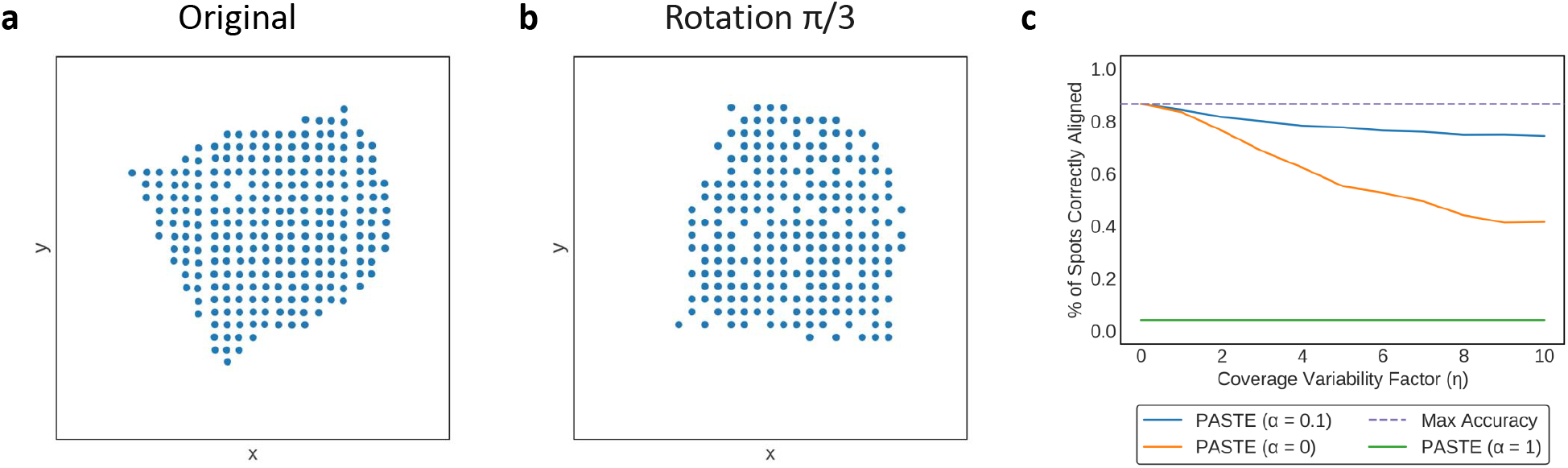
PASTE results for pairwise alignment of a simulated ST layer and an ST layer 1 from the breast cancer dataset from [42]. (a) Spatial organization of 254 spots from layer 1 of breast cancer dataset. (b) Spatial organization of 220 spots from simulated layer with a rotation of 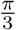. (c) Average percentage of spots correctly aligned by PASTE using *α* = 0 (gene expression data only), *α* = 1 (spatial information only), and *α* = 0.1 (both) as a function of the coverage variability factor *η*. Results are averaged over 10 simulations.

We first evaluated the performance of PASTE on the Pairwise Layer Alignment Problem by aligning a real breast cancer layer (*X, Z*) to simulated layers (*X′, Z′*) with a constant rotation of 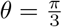 and increasing coverage variability factor *η* (Figure 2ab). Since the exact alignment of spots is known, we measured performance by computing ∑_*i~j*_ *π_ij_*, the percentage of spots correctly aligned between the original layer and simulated layer. We tested the performance of PASTE using only gene expression data (*α* = 0), using only spatial data (*α* = 1) and using both types of data (*α* = 0.1). When running PASTE, we used uniform distributions 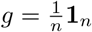 and 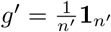, over spots in each layer.

We observe that PASTE achieves highest accuracy when using both gene expression and spatial information (*α* = 0.1), outperforming the alignment when using either expression or spatial information alone (Figure 2c and Supplementary Figure S3). Note that because the number of populated spots in each layer are not identical, perfect alignment corresponds to only 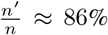 of spots being aligned. PASTE with *α* = 0.1 correctly aligns > 73% of spots even when the coverage variability factor is large. When using only spatial data (*α* = 1), optimal transport does not recover any matched pair of spots, demonstrating that the rotation used in the simulation perturbs the spatial data to a desirable degree. The mappings produced by PASTE are sparse and every spot in the first layers is mapped with non-zero coefficients to an average of 1.86 spots in the other layer (Supplementary Table S8). To demonstrate that PASTE’s use of both expression and spatial data in computing the alignment is beneficial, we also compared PASTE to applying optimal transport to integrated expression matrices using the single-cell RNA-seq integration method Scanorama [23] and found that PASTE had consistently higher accuracy (Supplementary Section S2.1).

Next, we evaluated the performance of PASTE on the Center Layer Integration Problem. First, we used the same simulation procedure described above to simulate three ST layers {(*X*^(*q*)^, *Z*^(*q*)^); *q* = 1, 2, 3}from a real ST experiment (*X, Z*). We simulate the gene expression information of each layer independently and simulate the spatial information by a rotation of either 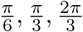 for each of the layers (Supplementary Figure S4). Since the exact alignment of spots from the center layer (*X, Z*) to each of the generated layers is known, we evaluated the performance of PASTE by computing 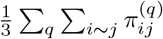, the average percentage of spots correctly aligned between the center layer and each of the three simulated layers. In addition, we compared the KL divergence between the gene expression matrix *X* of the true center layer and the low rank gene expression matrix *WH* inferred by PASTE.

We find that PASTE has both high accuracy and low difference reconstructing the true expression matrix. PASTE (*α* = 0.1) correctly aligns 49 – 54 % of spots (compared to maximum possible accuracy of 86%maximum accuracy) even with large values of *η*(Figure 3a and Supplementary Figure S5). In contrast, center layer integration using only gene expression data (*α* = 0) or using only spatial data (*α* = 1) performed poorly with accuracy below 2%. We also compared PASTE to the single cell RNAseq integration method Scanorama [23]. To do so, we ran Scanorama on the three simulated gene expression matrices of the ST layers to obtain 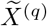, a batch corrected expression matrix for each layer q. Next, we compared the average difference between the true center expression matrix *X* and the corrected expression matrices 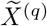 from Scanorama to the difference between the true expression matrix *X* and the integrated layer computed by PASTE; in both cases the difference between the matrices was computed using the KL divergence. We find that PASTE infers a center layer expression matrix that is much closer to truth than the integrated gene expression matrices from Scanorama (Figure 3b). Moreover, the results for Scanorama are actually an upper bound on performance since we used the true correspondence between layers when computing the KL divergence between spots. At the same time, Scanorama was not designed to utilize spatial data, and so the better performance shown by PASTE does not indicate a deficiency of Scanorama on the scRNA-seq integration problem for which is was designed. Finally, we tested our assumption that that the integrated gene expression matrix *X* is low rank. To that end, we ran PASTE without the use of NMF in each iteration of Algorithm 1 and set the inferred integrated matrix as 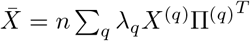. We see that PASTE with low rank assumption infers the center layer expression matrix more accurately than PASTE without NMF (Figure 3b and Supplementary Figure S6).

**Figure 3:**
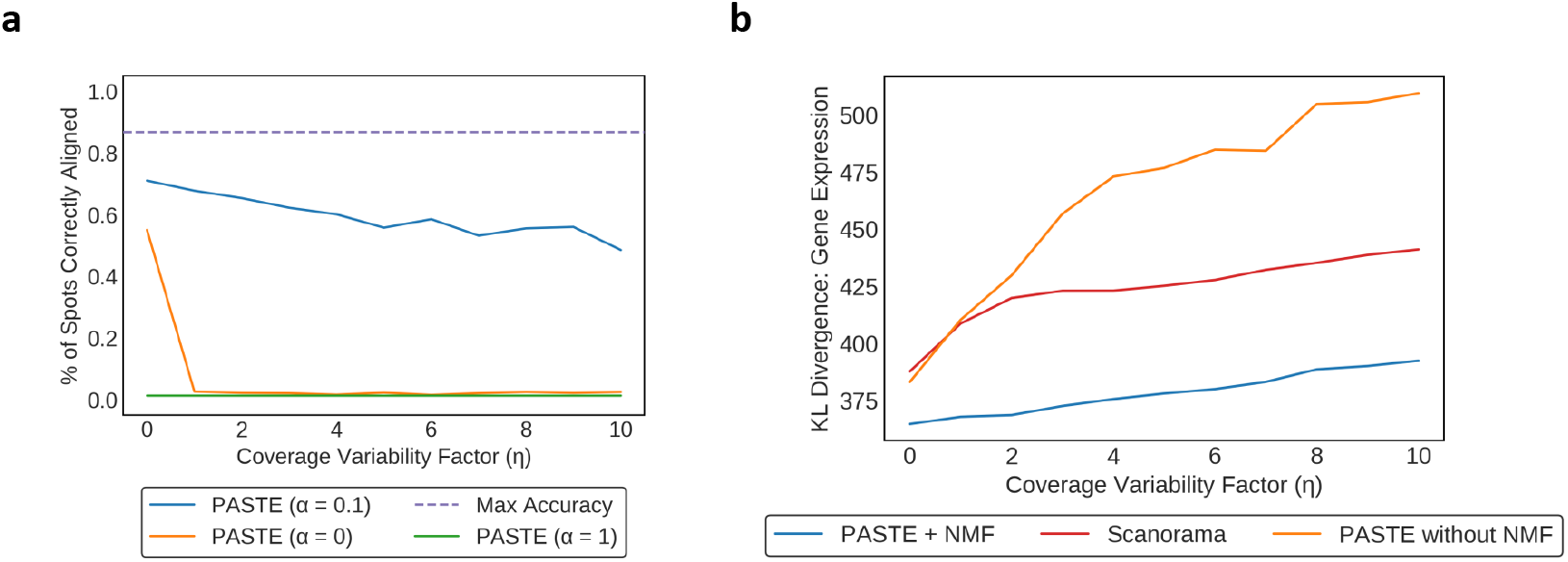
PASTE results on center layer alignment of simulated ST data from layer 2 of breast cancer dataset [42]. (a) Average percentage of spots correctly aligned between the original center layer and the simulated layers. (b) Difference between the gene expression matrix of the true center layer and the gene expression matrix inferred by PASTE and Scanorama. Differences are computed using KL divergence and for Scanorama, we computed the average KL divergence between the gene expression matrix of each of the three batch corrected simulated layers and the true center layer. PASTE without NMF dimensionality reduction was substantially worse than PASTE or Scanorama.

### 3.2 Analysis on Real Data

We applied PASTE to analyze ST datasets from four patients with cutaneous squamous cell carcinoma (SCC) [25]. Each patient in this dataset has three layers of ST data, with each layer containing ≈600 – 700 captured tissue spots. For each patient, [25] used independent component analysis to cluster the spots jointly across all three layers (Figure 4ab); this approaches uses only the gene expression data and does not utilize the spatial relationships between spots.

**Figure 4:**
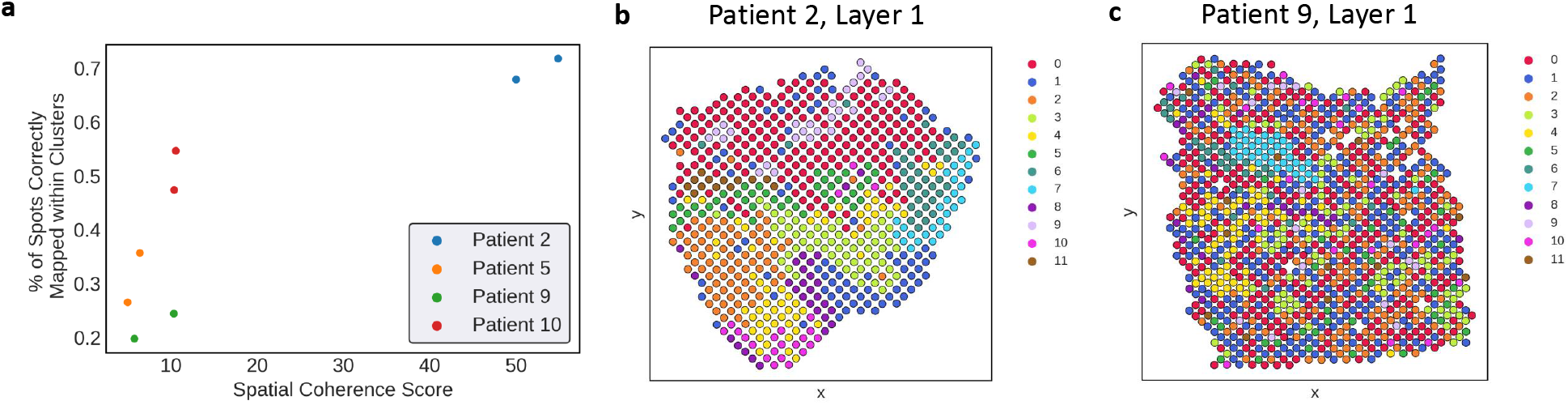
PASTE results on pairwise layer alignment of squamous cell carcinoma (SCC) ST dataset [25]. (a) Performance of PASTE in aligning spots from adjacent layers to same cluster as a function of the spatial coherence score. PASTE pairwise alignment shows greatest agreement with published cluster labels for patient 2 whose cluster labels also have the highest spatial coherence. (b) Published cluster labels of spots in layer 1 of patient 2 show moderate spatial coherence. (c) Published cluster labels of spots in layer 1 of patient 9 show low spatial coherence.

We first used PASTE to solve the Pairwise Layer Alignment Problem for adjacent tissue slices. The mappings produced by PASTE between pairs of adjacent layers were sparse having on average 1.7-2.1 mapped spots for each spot of the first layer (Supplementary Table S8). Under the assumption that spots adjacent to each other in 3D are likely to contain the same cell types (and thus the same expression cluster), we examined how frequently pairs of aligned spots determined by PASTE have the same cluster labels using the published cluster labels from [25]. Specifically, let *ℓ*(*i*) be the cluster labels of spot *i* and let Π be an alignment produced by PASTE between a pair of layers. We calculate ∑_*i,j*;*ℓ*(*i*)=*ℓ*(*j*)_ *π_ij_*: the fraction of aligned spots that have the same cluster labels in both layers.

We find that PASTE preserves cluster labels well in patient 2, but not as accurately in patients 5, 9, 10 (Figure 4a). For patient 2, around 70% of the spots in one layer are aligned to spots in the other layer with the same cluster labels. In contrast, for patients 5, 9, 10 only 20%-50% of aligned spots have the same cluster labels. There are multiple explanations for this result. First, it is possible that the published clusters for patients 5, 9, 10 are less reliable due to the fact that the sequence coverage for these patients was less than half of the sequence coverage in layers of patient 2. Second, it is possible that the published clusters and cluster labels are accurate, but there is less spatial structure in the tumors from patients 5, 9 and 10. Indeed there is a qualitative visual difference between the spatial coherence of clusters in patient 2 (Figure 4b) vs. the other patients (Figure 4c and Supplementary Figure S11). Since PASTE relies on spatial information to align layers, PASTE’s performance may decrease in datasets with less spatial coherence.

To quantify the observed differences in spatial coherence of clusters in different patients, we computed the *spatial entropy* of the cluster labels using O’Neil’s spatial entropy [37], which is the Shannon entropy of the observed distribution of pairs of cluster labels at adjacent spots in an ST layer (Supplementary Section S2.2). Since the range of spatial entropy values depends on both the number of clusters and the total number of spots, spatial entropy values are not directly comparable across patients or layers having different number of clusters and spots. Therefore, we compute a *Z*-score for spatial entropy by permuting cluster labels across the spots in a layer (Supplementary Figure S7), and define the *spatial coherence score* of a cluster labeling of a layer as the absolute value of the *Z*-score. This spatial coherence score gives a normalized quantity to compare spatial entropy values across patients and layers; a high spatial coherence score indicates that adjacent spots have the same cluster label more frequently than expected. Finally, we define the spatial coherence score for an aligned pair of layers to be the average spatial coherence score of the two layers.

We find that patient 2 has significantly higher spatial coherence scores than the other 3 patients, quantifying the observation that the cluster labels in patient 2 are the most spatially coherent (Figure 4a). While it is possible that the other three patients (5, 9 and 10) have less spatially coherent tumors, the fact that patients 5, 9 and 10 have less than half of the sequence coverage of patient 2 strongly suggests that the published clusters for these three patients may be problematic.

We constructed a 3D spatial representation of the SCC tumor from each patient using the pairwise alignments of adjacent layers computed by PASTE and projecting all layers to the same coordinate system by solving a generalized weighted Procrustes problem. We observe that PASTE recovers the 3D structure of the tumor despite the different placements and orientations of each tissue slice on the ST array (Figure 5 and Supplementary Figure S10). We further observe that the spatial distribution of clusters in patient 2 is indeed more coherent in 3D than the other patients.

**Figure 5:**
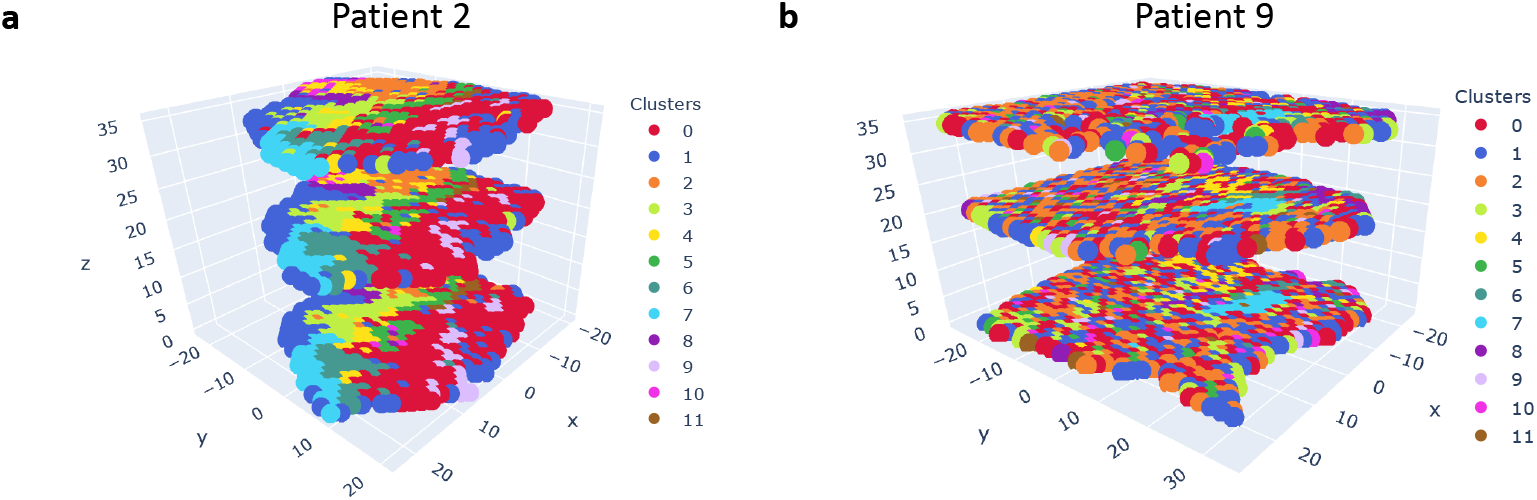
A 3D reconstruction of SCC tumors produced by PASTE using pairwise alignments of adjacent layers. Layers are colored according to published cluster labels [25]. (a) Aligned layers of patient 2. (b) Aligned layers of patient 9.

Next, we used PASTE to solve the Center Layer Integration Problem and infer a single center layer that integrate the multiple ST layers from each SCC patient. To evaluate the inferred center layer, we computed clusters from the inferred center layer expression matrix *X* = *WH* and evaluated the spatial coherence of these clusters, reasoning that expression clusters (representing cell types/states) should exhibit moderate spatial coherence in a tissue. Specifically, we applied *k*-means clustering to cluster spots according to the log normalized coordinates in the lower-dimensional space given by H. We set the number *k* of clusters equal to the published analysis of each patient [25].

We find that for all patients the clusters obtained using the center layer computed by PASTE have higher spatial coherence scores than the spatial coherence scores of the published clusters (Figure 6a and Supplementary Figure S11). Moreover, we see that the improvement in spatial coherence score is greatest in patients 5, 9, and 10 that have lower sequence coverage ST data. As one example, we note that while the the published cluster labels on layer 1 of patient 5 do not display much cluster coherence (Figure 6b), the cluster labels obtained from PASTE (Figure 6c) are visually much more spatially coherent. This shows that integrating multiple ST layers can recover subtle gene expression patterns which should improve downstream analysis of ST data.

**Figure 6:**
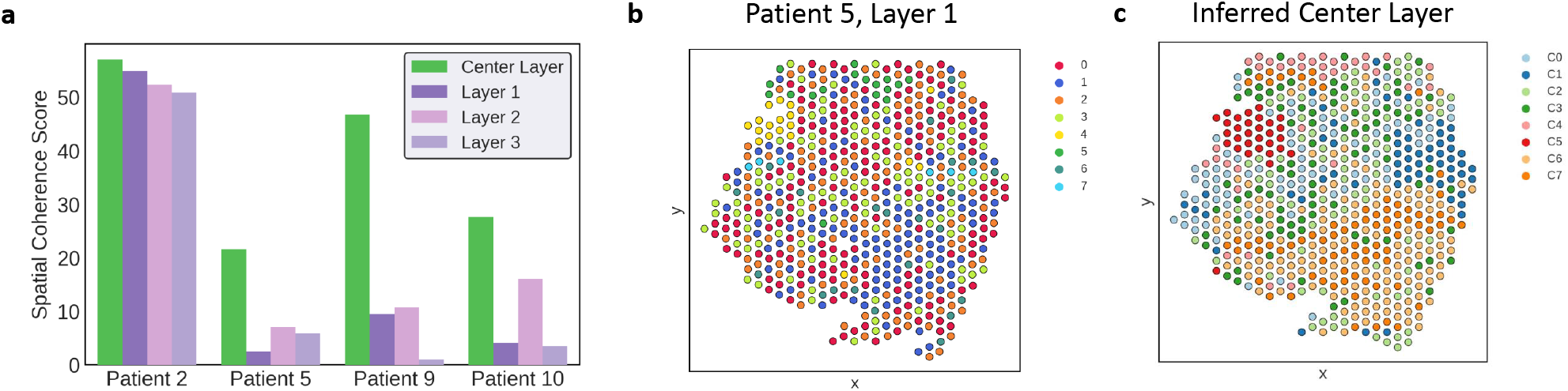
PASTE results on integration of multiple layers from SCC ST dataset [25] into a center layer. (a) Spatial coherence scores for the clusters obtained from the center layer inferred by PASTE (green) and the published clusters from [25] on the individual layers from each patient. The center layer inferred by PASTE has substantially high spatial coherence. (b) Published cluster labels of spots in layer 1 of patient 5. (c) Cluster labels *C*_1_,…, *C*_7_ of spots obtained from PASTE’s inferred center layer for patient 5.

Finally, we evaluated PASTE on an additional SCC patient from [25] that has two layers of ST data obtained using the newer, higher-resolution Visium platform [1]. The dataset has a higher number of spots per layer (722 and 674) and a higher number of transcripts per spot (16847 median) compared to the earlier ST platform used in patients 2, 5, 9, and 10. Consistent with the ST analysis of the first four patients, we find that the center layer inferred by PASTE gives more spatially coherent clusters than the clusters obtained on individual layers (Supplementary Figure S12). This demonstrates the ability of PASTE to scale to ST experiments from the latest ST technologies.

## 4 Discussion

In this paper, we introduce PASTE (Probabilistic Alignment of ST Experiments), a method to align and integrate replicate ST experiments by leveraging both transcriptional similarity and spatial distances between spots. We formalize and solve the Pairwise Layer Alignment Problem of mapping spots across adjacent ST layers and the Center Layer Integration Problem of integrating multiple ST layers by finding a center layer with a low rank expression matrix and mappings of its spots to all other layers. We show these problems can be solved using fused Gromov-Wasserstein optimal transport (FGW-OT) for the pairwise problem, and alternating between solving instances of the FGW-OT problem and performing NMF of a weighted expression matrix for the center problem. On simulated data, we show that PASTE’s use of both transcriptional and spatial data outperforms using either data modality alone. On real ST from squamous cell carcinoma, we show that the center layer inferred by PASTE has higher spatial coherence than published clusters that were inferred from ST data without considering spatial coordinates of spots. Interestingly, we see that while published clusters inferred from patients with high coverage ST data showed high spatial coherence, clusters inferred from patients with low coverage ST data had low spatial coherence. In comparison, clusters inferred on the center expression matrix inferred PASTE had high spatial coherence across all patients. Furthermore, by stacking adjacent layers one on top of the other we see that the spatial coherence within each layer is preserved also in 3D for patients with high sequencing coverage. We applied PASTE to data from ST technology and the newer Visium technology from 10X Genomes, but note that PASTE can also be applied to other spatial technologies such as smFISH [26], seqFISH+[18], and STARmap [51].

PASTE has some limitations and can be improved in several ways. First, in cases were the tissue is symmetrical, our alignment may be ambiguous since spots may be equally mapped to several locations. Second, our model assumes that each spot in one layer is represented as a convex combination of spots in the other layer, whereas the true mapping may in fact be non-linear. Third, our model does not use the histological images that often accompany the ST tissues in order to align and merge layers. In contrast, a recent software package, STUtility [4], aligns the histological images without considering the accompanying gene expression data. We anticipate PASTE could be further improved by utilizing the histological images and using methods from the realm of image registration [8].

The aligned and integrated ST layers produced by PASTE can be used to increase the statistical power in multiple downstream analyses including: identification of spatial expression patterns [42, 30], spatial cell type annotation [5], tumor/normal spot classification [52], spatial cell-cell communication patterns [2, 9], identification of genomic copy number aberrations [17], and more. In addition, PASTE could be applied to ST experiments from different patients in order to find conserved patterns of gene expression across different patients. Finally, newer versions of the Visium platform are now able to measure protein immunofluorescence in conjunction to gene expression, thus opening new analysis opportunities.

## 5 Acknowledgements

This work is supported by grants U24CA211000 and U24CA248453 from the US National Cancer Institute (NCI).

## Footnotes

1 Without loss of generality, we assume that a distribution *g_i_* is strictly positive, since spots *i* with *g_i_* = 0 can be removed.

## References

[1] 10x Genomics. Visium spatial gene expression: Map the whole transcriptome within the tissue context, 2019. Accessed: october 2020.

[2] Damien Arnol, Denis Schapiro, Bernd Bodenmiller, Julio Saez-Rodriguez, and Oliver Stegle. Modeling cell-cell interactions from spatial molecular data with spatial variance component analysis. Cell Reports, 29(1):202–211, 2019.

[3] Michaela Asp, Fredrik Salmén, Patrik L Ståhl, Sanja Vickovic, Ulrika Felldin, Marie Löfling, José Fernandez Navarro, Jonas Maaskola, Maria J Eriksson, Bengt Persson, et al. Spatial detection of fetal marker genes expressed at low level in adult human heart tissue. Scientific reports, 7(1):1–10, 2017.

[4] Joseph Bergenstråhle, Ludvig Larsson, and Joakim Lundeberg. Seamless integration of image and molecular analysis for spatial transcriptomics workflows. BMC Genomics, 21(1):482, 2020.

[5] Emelie Berglund, Jonas Maaskola, Niklas Schultz, Stefanie Friedrich, Maja Marklund, Joseph Bergenstråhle, Firas Tarish, Anna Tanoglidi, Sanja Vickovic, Ludvig Larsson, Fredrik Salmén, Christoph Ogris, Karolina Wallenborg, Jens Lagergren, Patrik Ståhl, Erik Sonnhammer, Thomas Helleday, and Joakim Lundeberg. Spatial maps of prostate cancer transcriptomes reveal an unexplored landscape of heterogeneity. Nature Communications, 9(1):2419, 2018.

[6] Garrett Birkhoff. Tres observaciones sobre el algebra lineal. Univ. Nac. Tucuman, Ser. A, 5:147–154, 1946.

[7] Vincent D Blondel, Ngoc-Diep Ho, Paul Dooren, et al. Weighted nonnegative matrix factorization and face feature extraction. In In Image and Vision Computing. Citeseer, 2008.

[8] Lisa Gottesfeld Brown. A survey of image registration techniques. ACM Comput. Surv., 24(4):325–376, December 1992.

[9] Zixuan Cang and Qing Nie. Inferring spatial and signaling relationships between cells from single cell transcriptomic data. Nature Communications, 11(1):2084, 2020.

[10] Wei-Ting Chen, Ashley Lu, Katleen Craessaerts, Benjamin Pavie, Carlo Sala Frigerio, Nikky Corthout, Xiaoyan Qian, Jana Laláková, Malte Kühnemund, Iryna Voytyuk, Leen Wolfs, Renzo Mancuso, Evgenia Salta, Sriram Balusu, An Snellinx, Sebastian Munck, Aleksandra Jurek, Jose Fernandez Navarro, Takaomi C. Saido, Inge Huitinga, Joakim Lundeberg, Mark Fiers, and Bart De Strooper. Spatial transcriptomics and *in situ* sequencing to study alzheimer’s disease. Cell, 182(4):976–991.e19, 2020/10/26 2020.

[11] Marco Cuturi. Sinkhorn distances: Lightspeed computation of optimal transport. In C. J. C. Burges, L. Bottou, M. Welling, Z. Ghahramani, and K. Q. Weinberger, editors, Advances in Neural Information Processing Systems, volume 26, pages 2292–2300. Curran Associates, Inc., 2013.

[12] Marco Cuturi and Arnaud Doucet. Fast computation of wasserstein barycenters. volume 32 of Proceedings of Machine Learning Research, pages 685–693, Bejing, China, 22-24 Jun 2014. PMLR.

[13] Pinar Demetci, Rebecca Santorella, Björn Sandstede, William Stafford Noble, and Ritambhara Singh. Gromov-wasserstein optimal transport to align single-cell multi-omics data. bioRxiv, 2020.

[14] Ghislain Durif, Laurent Modolo, Jeff E Mold, Sophie Lambert-Lacroix, and Franck Picard. Probabilistic count matrix factorization for single cell expression data analysis. Bioinformatics, 35(20):4011–4019, 03 2019.

[15] Marc Elosua, Paula Nieto, Elisabetta Mereu, Ivo Gut, and Holger Heyn. Spotlight: Seeded nmf regression to deconvolute spatial transcriptomics spots with single-cell transcriptomes. bioRxiv, 2020.

[16] Rebecca Elyanow, Bianca Dumitrascu, Barbara E. Engelhardt, and Benjamin J. Raphael. netnmf-sc: leveraging gene–gene interactions for imputation and dimensionality reduction in single-cell expression analysis. Genome Research, 30(2):195–204, 2020.

[17] Rebecca Elyanow, Ron Zeira, Max Land, and Benjamin Raphael. STARCH: Copy number and clone inference from spatial transcriptomics data. Physical Biology, oct 2020.

[18] Chee-Huat Linus Eng, Michael Lawson, Qian Zhu, Ruben Dries, Noushin Koulena, Yodai Takei, Jina Yun, Christopher Cronin, Christoph Karp, Guo-Cheng Yuan, et al. Transcriptome-scale superresolved imaging in tissues by rna seqfish+. Nature, 568(7751):235–239, 2019.

[19] Rémi Flamary and Nicolas Courty. Pot python optimal transport library, 2017.

[20] C. Févotte and A. T. Cemgil. Nonnegative matrix factorizations as probabilistic inference in composite models. In 2009 17th European Signal Processing Conference, pages 1913–1917, 2009.

[21] Dan Gusfield. Algorithms on stings, trees, and sequences: Computer science and computational biology. Acm Sigact News, 28(4):41–60, 1997.

[22] Laleh Haghverdi, Aaron TL Lun, Michael D Morgan, and John C Marioni. Batch effects in single-cell rna-sequencing data are corrected by matching mutual nearest neighbors. Nature biotechnology, 36(5):421–427, 2018.

[23] Brian Hie, Bryan Bryson, and Bonnie Berger. Efficient integration of heterogeneous single-cell transcriptomes using scanorama. Nature Biotechnology, 37(6):685–691, 2019.

[24] Wenpin Hou, Zhicheng Ji, Hongkai Ji, and Stephanie C. Hicks. A systematic evaluation of singlecell rna-sequencing imputation methods. Genome Biology, 21(1):218, 2020.

[25] Andrew Ji, Adam Rubin, Kim Thrane, Sizun Jiang, David Reynolds, Robin Meyers, Margaret Guo, Benson George, Annelie Mollbrink, Joseph Bergenstråhle, Ludvig Larsson, Yunhao Bai, Bokai Zhu, Aparna Bhaduri, Jordan Meyers, Xavier Rovira-Clavé, S Hollmig, Sumaira Aasi, Garry Nolan, and Paul Khavari. Multimodal analysis of composition and spatial architecture in human squamous cell carcinoma. Cell, 182:1661–1662, 09 2020.

[26] Ni Ji and Alexander Oudenaarden. Single molecule fluorescent in situ hybridization (smfish) of c. elegans worms and embryos. WormBook : the online review of C. elegans biology, pages 1–16, 12 2012.

[27] W. Kabsch. A solution for the best rotation to relate two sets of vectors. Acta Crystallographica Section A, 32(5):922–923, Sep 1976.

[28] Daniel D. Lee and Hyunjune Sebastian Seung. Algorithms for non-negative matrix factorization. In Advances in Neural Information Processing Systems 13 - Proceedings of the 2000 Conference, NIPS 2000, Advances in Neural Information Processing Systems. Neural information processing systems foundation, January 2001. 14th Annual Neural Information Processing Systems Conference, NIPS 2000; Conference date: 27-11-2000 Through 02-12-2000.

[29] Peijie Lin, Michael Troup, and Joshua W. K. Ho. Cidr: Ultrafast and accurate clustering through imputation for single-cell rna-seq data. Genome Biology, 18(1): 59, 2017.

[30] Ruishan Liu, Marco Mignardi, Robert Jones, Martin Enge, Seung K Kim, Stephen R Quake, and James Zou. Modeling spatial correlation of transcripts with application to developing pancreas. Scientific reports, 9(1):1–8, 2019.

[31] Anna Lundmark, Natalija Gerasimcik, Tove Båge, Anders Jemt, Annelie Mollbrink, Fredrik Salmén, Joakim Lundeberg, and Tülay Yucel-Lindberg. Gene expression profiling of periodontitis-affected gingival tissue by spatial transcriptomics. Scientific reports, 8(1):1–9, 2018.

[32] Igor Mandric, Brian L. Hill, Malika K. Freund, Michael Thompson, and Eran Halperin. Batman: Fast and accurate integration of single-cell rna-seq datasets via minimum-weight matching. iScience, 23(6):101185, 2020.

[33] Silas Maniatis, Tarmo Äijö, Sanja Vickovic, Catherine Braine, Kristy Kang, Annelie Mollbrink, Delphine Fagegaltier, Žaneta Andrusivová, Sami Saarenpää, Gonzalo Saiz-Castro, et al. Spatiotemporal dynamics of molecular pathology in amyotrophic lateral sclerosis. Science, 364(6435):89–93, 2019.

[34] Reuben Moncada, Dalia Barkley, Florian Wagner, Marta Chiodin, Joseph C. Devlin, Maayan Baron, Cristina H. Hajdu, Diane M. Simeone, and Itai Yanai. Integrating microarray-based spatial transcriptomics and single-cell rna-seq reveals tissue architecture in pancreatic ductal adenocarcinomas. Nature Biotechnology, 38(3):333–342, 2020.

[35] Aanchal Mongia, Debarka Sengupta, and Angshul Majumdar. Mcimpute: Matrix completion based imputation for single cell rna-seq data. Frontiers in Genetics, 10:9, 2019.

[36] Mor Nitzan, Nikos Karaiskos, Nir Friedman, and Nikolaus Rajewsky. Gene expression cartography. Nature, 576(7785):132–137, 2019.

[37] RV o’neill, JR Krummel, RH e al Gardner, G Sugihara, B Jackson, DL DeAngelis, BT Milne, Monica G Turner, B Zygmunt, SW Christensen, et al. Indices of landscape pattern. Landscape ecology, 1(3):153–162, 1988.

[38] Gabriel Peyré and Marco Cuturi. Computational optimal transport: With applications to data science. Foundations and Trends^®^ in Machine Learning, 11(5-6):355–607, 2019.

[39] Gabriel Peyré, Marco Cuturi, and Justin Solomon. Gromov-wasserstein averaging of kernel and distance matrices. volume 48 of Proceedings of Machine Learning Research, pages 2664–2672, New York, New York, USA, 20–22 Jun 2016. PMLR.

[40] Geoffrey Schiebinger, Jian Shu, Marcin Tabaka, Brian Cleary, Vidya Subramanian, Aryeh Solomon, Joshua Gould, Siyan Liu, Stacie Lin, Peter Berube, et al. Optimal-transport analysis of single-cell gene expression identifies developmental trajectories in reprogramming. Cell, 176(4):928–943, 2019.

[41] Chunxuan Shao and Thomas Höfer. Robust classification of single-cell transcriptome data by nonnegative matrix factorization. Bioinformatics, 33(2):235–242, 09 2016.

[42] Patrik L Ståhl, Fredrik Salmén, Sanja Vickovic, Anna Lundmark, José Fernández Navarro, Jens Magnusson, Stefania Giacomello, Michaela Asp, Jakub O Westholm, Mikael Huss, et al. Visualization and analysis of gene expression in tissue sections by spatial transcriptomics. Science, 353(6294):78–82, 2016.

[43] Tim Stuart, Andrew Butler, Paul Hoffman, Christoph Hafemeister, Efthymia Papalexi, III Mauck, William M., Yuhan Hao, Marlon Stoeckius, Peter Smibert, and Rahul Satija. Comprehensive integration of single-cell data. Cell, 177(7):1888–1902.e21, 2020/10/26 2019.

[44] Valentine Svensson, Sarah A Teichmann, and Oliver Stegle. Spatialde: identification of spatially variable genes. Nature methods, 15(5):343, 2018.

[45] Kim Thrane, Hanna Eriksson, Jonas Maaskola, Johan Hansson, and Joakim Lundeberg. Spatially resolved transcriptomics enables dissection of genetic heterogeneity in stage iii cutaneous malignant melanoma. Cancer research, 78(20):5970–5979, 2018.

[46] Vayer Titouan, Nicolas Courty, Romain Tavenard, and Rémi Flamary. Optimal transport for structured data with application on graphs. In International Conference on Machine Learning, pages 6275–6284, 2019.

[47] F. William Townes, Stephanie C. Hicks, Martin J. Aryee, and Rafael A. Irizarry. Feature selection and dimension reduction for single-cell rna-seq based on a multinomial model. Genome Biology, 20(1):295, 2019.

[48] Hoa Thi Nhu Tran, Kok Siong Ang, Marion Chevrier, Xiaomeng Zhang, Nicole Yee Shin Lee, Michelle Goh, and Jinmiao Chen. A benchmark of batch-effect correction methods for single-cell rna sequencing data. Genome Biology, 21(1): 12, 2020.

[49] Cédric Villani. Optimal transport: old and new, volume 338. Springer Science & Business Media, 2008.

[50] Grace Wahba. A least squares estimate of satellite attitude. SIAM Review, 7(3):409–409, 1965.

[51] Xiao Wang, William E Allen, Matthew A Wright, Emily L Sylwestrak, Nikolay Samusik, Sam Vesuna, Kathryn Evans, Cindy Liu, Charu Ramakrishnan, Jia Liu, et al. Three-dimensional intact-tissue sequencing of single-cell transcriptional states. Science, 361(6400), 2018.

[52] Niyaz Yoosuf, JoséFernández Navarro, Fredrik Salmén, Patrik L. Ståhl, and Carsten O. Daub. Identification and transfer of spatial transcriptomics signatures for cancer diagnosis. Breast Cancer Research, 22(1): 6, 2020.

[53] Xun Zhu, Travers Ching, Xinghua Pan, Sherman M. Weissman, and Lana Garmire. Detecting heterogeneity in single-cell rna-seq data by non-negative matrix factorization. PeerJ, 5: e2888, January 2017.

