## Supplement for "Alignment and Integration of Spatial Transcriptomics Data"

#### S1 Supplementary methods

##### S1.1 Proof of Theorem 1

**Theorem 1.** *Let* $\bar{X} = \sum_q \lambda_q X^{(q)} {\Pi^{(q)}}^T \operatorname{diag}(\frac{1}{g})$ *and* $X = WH$. *We have,*

$$S(W, H) = \sum_i g_i c(x_{\cdot i}, \bar{x}_{\cdot i}) + \tau$$

*where* $c(u, v) = \|u - v\|^2$ *or* $c(u, v) = KL(v||u) = \sum_l v_l \log \frac{v_l}{u_l} - v_l + u_l$, *and* $\tau$ *is a constant that does not depend on* $W, H$

*Proof.* We first prove the theorem for the Euclidean distance $c(u, v) = \|u - v\|^2$. We write the objective function explicitly and simplify it using $\sum_j \Pi_{ij}^{(q)} = g_i$ and $\sum_q \lambda_q = 1$.

$$\begin{aligned}
S(W, H) &= \sum_q \lambda_q \sum_i \sum_j \left\| x_{\cdot i} - x_{\cdot j}^{(q)} \right\|^2 \pi_{ij}^{(q)} \\
&= \sum_q \lambda_q \sum_i \sum_j {x_{\cdot i}}^T x_{\cdot i} \pi_{ij}^{(q)} - 2{x_{\cdot i}}^T x_{\cdot j}^{(q)} \pi_{ij}^{(q)} + \beta \\
&= \sum_i {x_{\cdot i}}^T x_{\cdot i} \sum_j \pi_{ij}^{(q)} \sum_q \lambda_q - 2 \sum_i \sum_j {x_{\cdot i}}^T \sum_q \lambda_q x_{\cdot j}^{(q)} \pi_{ij}^{(q)} + \beta \\
&= \sum_i g_i {x_{\cdot i}}^T x_{\cdot i} - 2 \operatorname{Tr}(X^T \sum_q \lambda_q X^{(q)} {\Pi^{(q)}}^T) + \beta \\
&= \operatorname{Tr}(X^T X \operatorname{diag}(g)) - 2 \operatorname{Tr}(X^T \sum_q \lambda_q X^{(q)} {\Pi^{(q)}}^T \operatorname{diag}(\frac{1}{g}) \operatorname{diag}(g)) + \beta \\
&= \operatorname{Tr}\big((X^T X - 2X^T \bar{X}\big) \operatorname{diag}(g)) + \beta \\
&= \sum_i g_i \left\| x_{\cdot i} - \bar{x}_{\cdot i} \right\|^2 + \beta'
\end{aligned}$$

where $\beta$ and $\beta'$ are constants that do not depend on $W, H$.

Next, we prove the theorem for the KL divergence $c(u,v) = KL(v||u) = \sum_l v_l \log \frac{v_l}{u_l} - v_l + u_l$. Again, we write the objective function explicitly and simplify it:

$$\begin{aligned} S(W,H) &= \sum_q \lambda_q \sum_i \sum_j \Big[ \sum_l x_{li} - x_{lj}^{(q)} - x_{lj}^{(q)} \log(x_{li}) + x_{lj}^{(q)} \log(x_{lj}^{(q)}) \Big] \pi_{ij}^{(q)} \\ &= \sum_q \lambda_q \sum_i \sum_j \Big[ \sum_l x_{li} - x_{lj}^{(q)} \log(x_{li}) \Big] \pi_{ij}^{(q)} + \gamma \\ &= \sum_i \sum_l \Big[ x_{li} \sum_j \pi_{ij}^{(q)} \sum_q \lambda_q - \log(x_{li}) \sum_j \sum_q \lambda_q x_{lj}^{(q)} \pi_{ij}^{(q)} \Big] + \gamma \\ &= \sum_i g_i \sum_l \Big[ x_{li} - \bar{x}_{li} \log(x_{li}) \Big] + \gamma \\ &= \sum_i g_i \mathrm{KL}(\bar{x}_{\cdot i} || x_{\cdot i}) + \gamma' \end{aligned}$$

where $\gamma$ and $\gamma'$ are constants that do not depend on $W, H$. □

#### S1.2 Finding optimal rotation for spatial coordiantes

In this section, we seek to find a rotation and translation that of the spatial coordinates of one layer that minimizes the distances to the spatial coordinates of the other layer given a mapping. The problem of finding rotation and translation that minimizes the distances between matched set of points is a well know problem in several research fields [6, 3]. In 2d the problem is often called called Procrustes analysis , a more general linear algebra problem is called the Orthogonal Procrustes problem , and the vector weighted version is called Wahba's problem [6]. In chemistry/biology the solution to the 3d problem is called the Kabsch algorithm [3]. The 2d solution is based on finding the rotation angle while the general case (which also works in 2d) looks for a rotation matrix, thus it also supports reflection.

Our problem is a variation of this problem since we have a probabilistic alignment between the spots given by the mapping $\Pi$.

**Problem S1.** *Given ST layers with spatial coordinates $Z \in \mathbb{R}^{2\times n}$ and $W \in \mathbb{R}^{2\times n'}$ and a mapping $\Pi \in \Gamma(g, g')$, find a vector $t \in \mathbb{R}^2$ and a rotation matrix $R \in \mathbb{R}^{2\times 2}$:*

$$Q(t,R) = \sum_{i,j} \pi_{ij} \left\| z_{\cdot i} - R w_{\cdot j} - t \right\|^2. \tag{S1}$$

We first show that we can assume that no translation is needed ($t = 0$) by centering the spatial coordinates $Z$ and $W$. Assuming $R$ is fixed, we can find the optimal translation by taking the derivative of $Q$ w.r.t. $t$ and comparing to zero:

$$
\begin{aligned}
\frac{\partial Q}{\partial t} &= -2\sum_{i,j}\pi_{ij}(z_{\cdot i} - Rw_{\cdot j} - t) \\
&= -2\sum_{i} z_{\cdot i}\sum_{j}\pi_{ij} + 2\sum_{j} w_{\cdot j}\sum_{i}\pi_{ij} + 2t\sum_{i,j}\pi_{ij} \\
&= -2\sum_{i} z_{\cdot i}g_i + 2\sum_{j} w_{\cdot j}g'_j + 2t = 0
\end{aligned}
$$

We have $\hat{t} = Zg - Wg'$. By replacing the spatial coordinates $z_{\cdot i}$ with $z_{\cdot i} - Zg$ and the spatial coordinates $w_{\cdot j}$ with $w_{\cdot j} - Wg'$ we get $Q = \sum_{i,j}\pi_{ij}\left\|z_{\cdot i} - Rw_{\cdot j}\right\|^2$. Therefore, centering both spatial coordinates removes the need to find a translation and we are only left with finding the optimal rotation.

We rewrite the objective $Q$ in matrix notation:

$$
\begin{aligned}
Q &= \sum_{i,j}\pi_{ij}(z_{\cdot i} - Rw_{\cdot j}) \\
&= \sum_{i,j}\pi_{ij}(z_{\cdot i}^T z_{\cdot i} - w_{\cdot j}^T R^T R w_{\cdot j} - z_{\cdot i}^T R w_{\cdot j} - w_{\cdot j}^T R^T z_{\cdot i}) \\
&= -2\sum_{i,j}\pi_{ij}(z_{\cdot i}^T - Rw_{\cdot j}) + \alpha \\
&= -2\operatorname{Tr}(Z^T R W \Pi^T) + \alpha \\
&= -2\operatorname{Tr}(R W \Pi^T Z^T) + \alpha
\end{aligned}
$$

where $\alpha$ us a constant independent of $R$.

We find the optimal rotation $R$ that minimizes $Q$ using SVD similar to the solution to Wahbs's problem [4]. Let $U\Sigma V^T$ be the SVD decomposition of $W\Pi^T Z^T$. We have

$$
\begin{aligned}
Q &= -2\operatorname{Tr}(RU\Sigma V^T) + \alpha \\
&= -2\operatorname{Tr}(\Sigma V^T R U) + \alpha
\end{aligned}
$$

Notice that $\Sigma$ is a positive diagonal matrix and $V^T RU$ is an orthonormal matrix. Therefore, the objective $Q$ is minimized when the trace of $V^T RU$ is maximal which is attained when $V^T RU = I$. We have $R = VU^T$. We note that $R$ may also do reflection in addition to rotation.

An alternative derivation for the 2d case is done similar to Procrustes analysis. We write the rotation matrix as a function of the rotation angle $\theta$:

$$
R(\theta) = \begin{pmatrix} \cos(\theta) & -\sin(\theta) \\ \sin(\theta) & \cos(\theta) \end{pmatrix}
$$

Taking the derivative of $Q$ with respect to $\theta$ and comparing to zero gives:

$$
\begin{aligned}
\frac{\partial Q}{\partial \theta} &= -2\,\mathrm{Tr}\Big(\frac{\partial R(\theta)}{\partial \theta} W\Pi^T Z^T\Big) \\
&= -2\,\mathrm{Tr}\Big(\begin{pmatrix} -\sin(\theta) & -\cos(\theta) \\ \cos(\theta) & -\sin(\theta)\end{pmatrix} W\Pi^T Z^T\Big) = 0
\end{aligned}
$$

Dividing by $\cos(\theta)$ and extracting $\theta$ we have:

$$
\hat{\theta} = \arctan\Big(\frac{\mathrm{Tr}\Big(\begin{pmatrix} 0 & -1 \\ 1 & 0\end{pmatrix} W\Pi^T Z^T\Big)}{\mathrm{Tr}(W\Pi^T Z^T)}\Big)
$$

### S2 Supplementary results

#### S2.1 Comparison of PASTE to Scanorama on ST alignment simulation

We compared our results to a SC-RNAseq integration method Scanorama [1]. Scanorama integrates gene expression information by resolving noise and batch effects between two or more datasets. Scanorama is not designed to align cells from RNAseq, though it does relies on inferring nearest neighbors between cells in the given data sets. To directly compare Scanorama with PASTE, we calculated an alignment between spots of the different layers by finding a mapping that minimizes the Wasserstein optimal transport distance, where the transportation cost between the spots is taken as the Euclidean distance between the spots in the integrated gene expression datasets from Scanorama.

We see that PASTE outperforms alignment based gene expression corrected by Scanorama (Figure S3). In fact, Scanorama performs slightly worse than our pairwise alignment on the original gene expression data alone.

#### S2.2 Spatial entropy definition

The spatial entropy is computed as follows. Given a graph with vertex labels (e.g. cluster labels), the spatial entropy is the Shannon entropy of the distribution of the unordered pairs of cluster labels on the edges of the graph. Specifically, let $G = (V, E)$ be graph where $V$ is the set of spots and where there is an edge $(i, k) \in E$ between every pair $(i, k)$ of spots adjacent on the array. Let $K = \{1, 2, \ldots, k\}$ be a set of $k$ cluster labels and let $\ell : V \leftarrow K$ be the spot cluster assignment. We define a categorical variable $C = \{\{a, b\}; (a, b) \in N \times N\}$ which describes every distinct unordered pair of cluster labels. The spatial entropy is calculated as $H(G) = H(C|E) = -\sum_{c\in C} \mathbb{P}(c|E)\log(\mathbb{P}(c|E))$, where $\mathbb{P}(c|E) = \frac{c}{|E|}$. A low value of spatial entropy indicates that many adjacent spots have the same cluster label, while a large spatial entropy indicates that many adjacent spots have different cluster labels.

#### S2.3 Supplementary plots

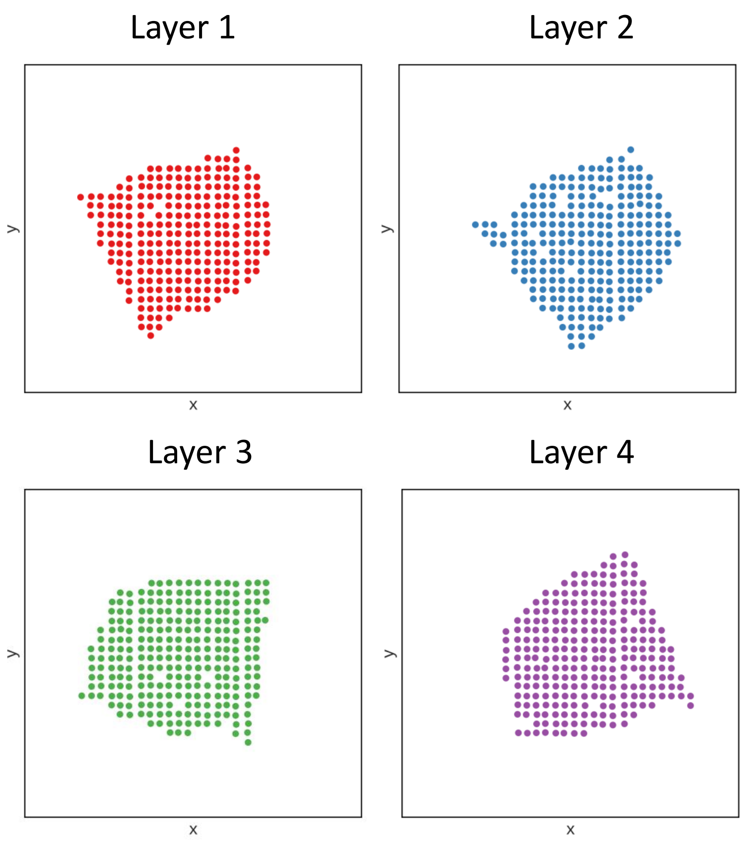


Figure S1: Spatial organization of breast cancer ST layers from [5].

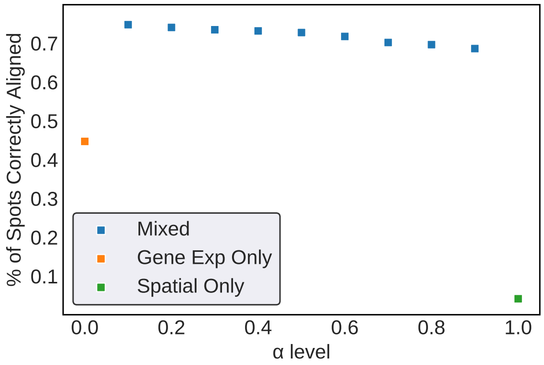


Figure S2: PASTE results for pairwise alignment of a simulated ST layer with layer 1 of breast cancer dataset [5] with varying levels of $\alpha$. Coverage variability factor for the simulated ST layer was set at $\eta = 10$.

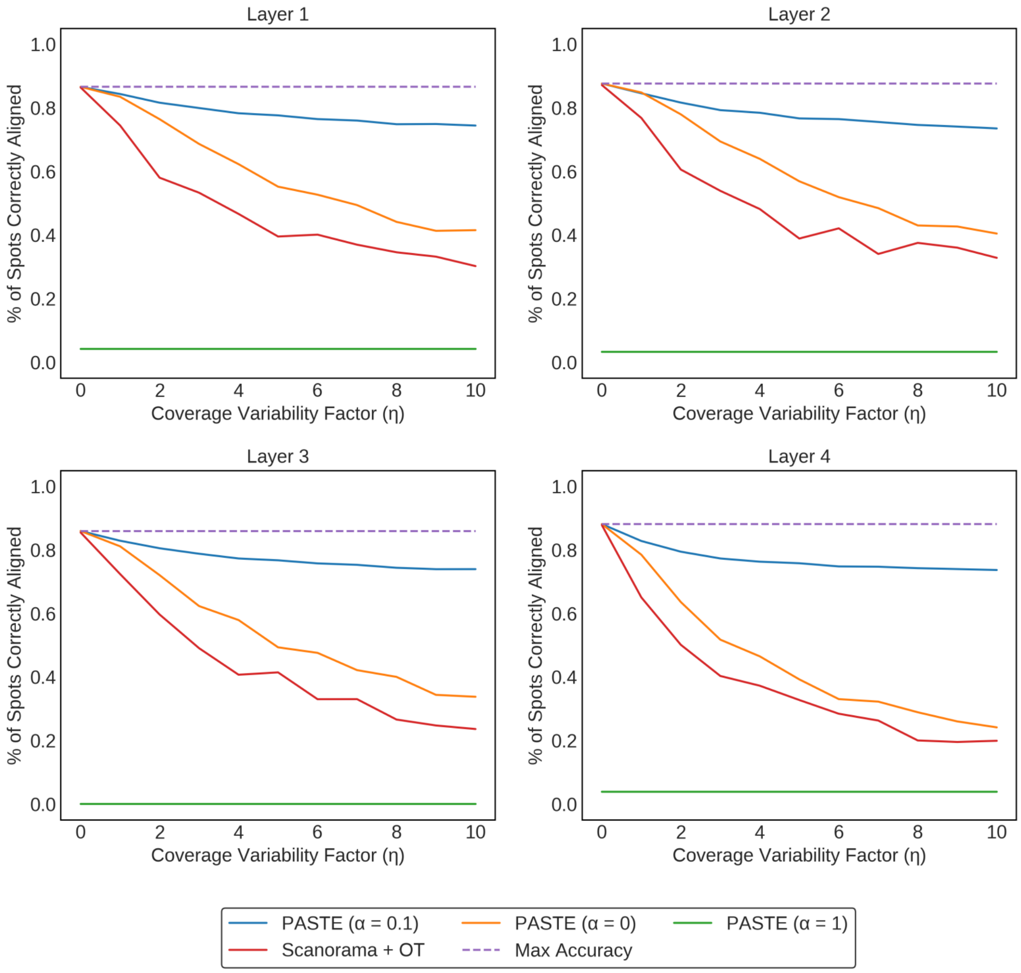


Figure S3: PASTE results on pairwise alignment of simulated ST layers based on four layers of breast cancer dataset [5]. Each value is an averaged over 10 simulations.

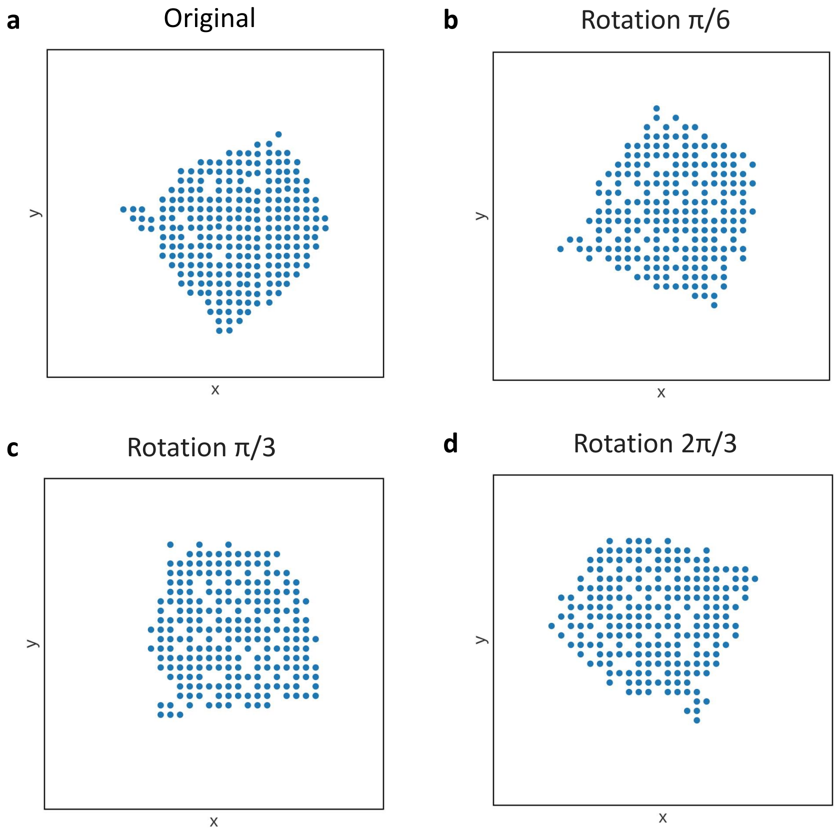


Figure S4: Spatial organization of spots used in center layer alignment simulation of layer 2 from the breast cancer dataset [5]. (a) Original spatial organization of spots in layer 2 of breast cancer dataset. (b) - (d) Simulated spatial structures obtained by rotating (a) by $\frac{\pi}{6}$, $\frac{\pi}{3}$, $\frac{2\pi}{3}$ respectively.

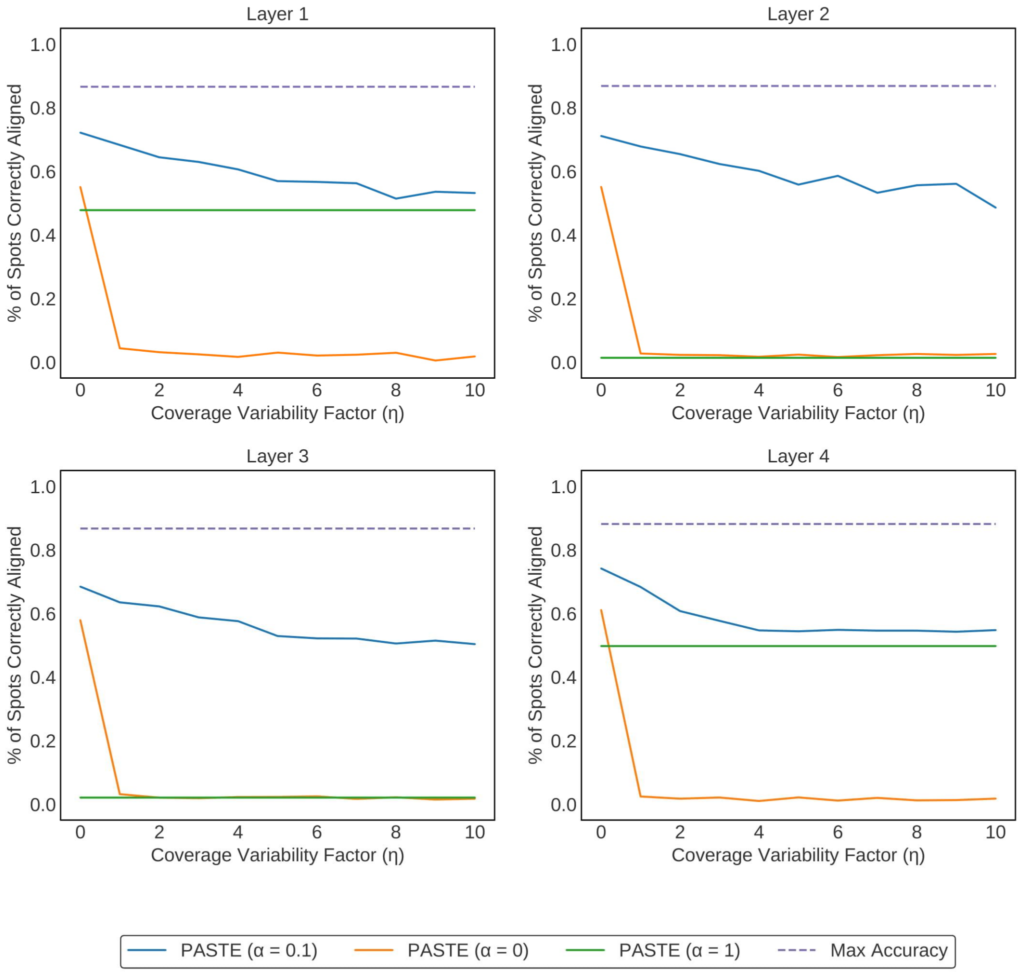


Figure S5: PASTE results on center layer integration of simulated ST layers based on four layers of breast cancer dataset [5].

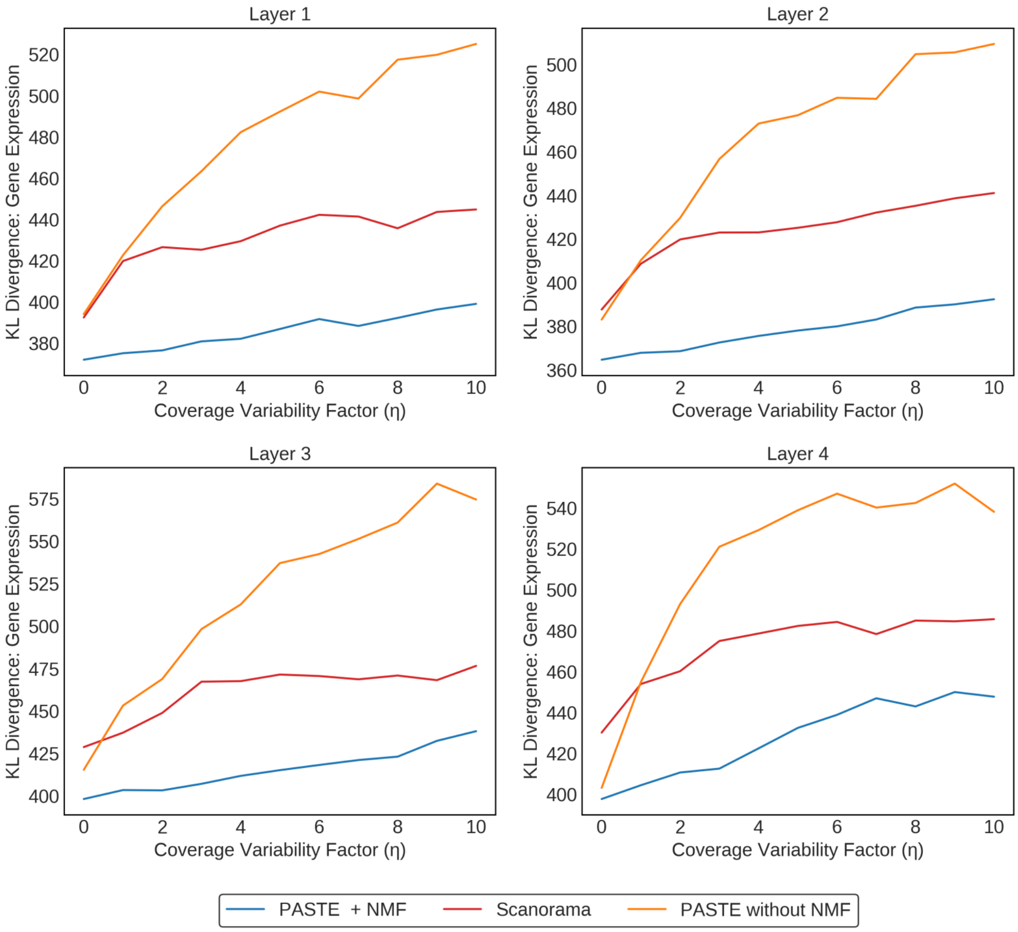


Figure S6: PASTE results on center layer integration of simulated ST layers compared to Scanorama and PASTE without NMF based on four layers of breast cancer dataset [5]. For this simulation, we used a pseudocount = 1.

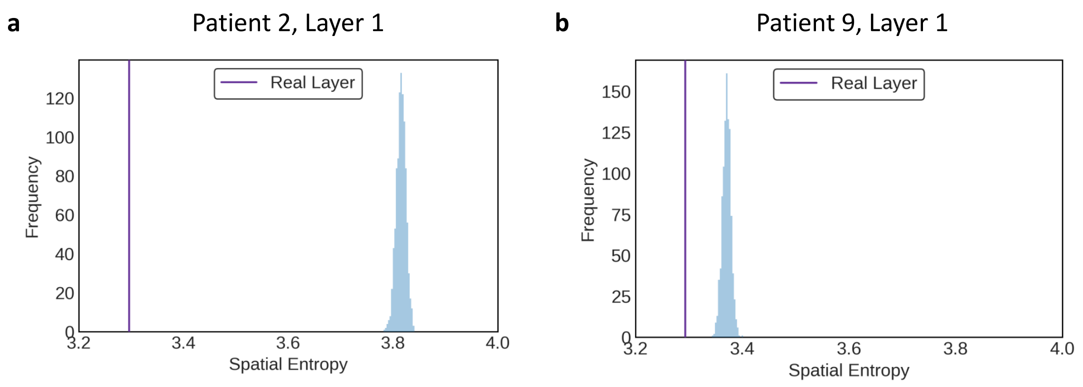


Figure S7: Histogram of spatial entropies. Given the cluster assignments on a real layer, we calculated the spatial entropy for 1000 random permutations of cluster labels on the spots. This distribution was used to calculate a spatial entropy z-score for the real layer. (a) Histogram of spatial entropies for patient 2, layer 1. (b) Histogram of spatial for patient 9, layer 1.

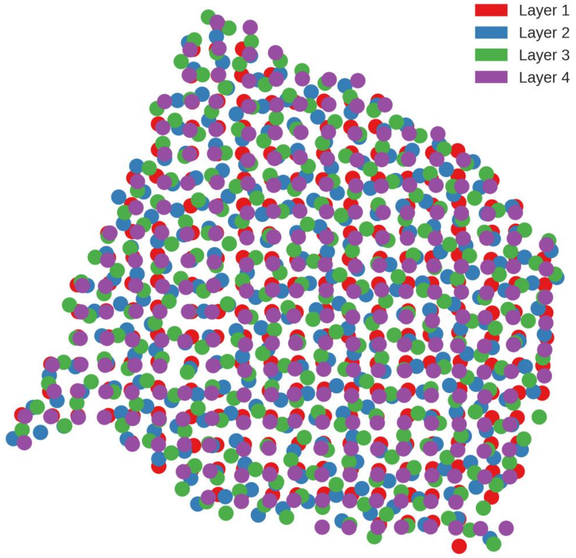


Figure S9: Spatial coordinates of the four breast cancer ST layers from [5] after pairwise alignment via PASTE.

**a** **Simulation Pairwise**

| | Layer 1 | Layer 2 | Layer 3 | Layer 4 |
|---|---|---|---|---|
| **Mixed** | 1.862 | 1.872 | 1.856 | 1.877 |
| **Gene Exp Only** | 1.862 | 1.872 | 1.856 | 1.877 |
| **Spatial Only** | 1.862 | 1.872 | 1.856 | 1.877 |

**b** **Simulation Center**

| | Layer 1 | Layer 2 | Layer 3 | Layer 4 |
|---|---|---|---|---|
| **Center, Layer 1** | 1.862 | 1.860 | 1.867 | 1.877 |
| **Center, Layer 2** | 1.862 | 1.872 | 1.856 | 1.877 |
| **Center, Layer 3** | 1 | 1 | 1 | 1 |

**c** **SCC Pairwise**

| | Patient 2 | Patient 5 | Patient 9 | Patient 10 |
|---|---|---|---|---|
| **Layers 1, 2** | 1.968 | 1.88 | 1.934 | 2.019 |
| **Layers 2, 3** | 1.986 | 1 | 2.102 | 1.742 |

**d** **SCC Center**

| | Patient 2 | Patient 5 | Patient 9 | Patient 10 |
|---|---|---|---|---|
| **Center, Layer 1** | 1 | 1 | 1 | 1 |
| **Center, Layer 2** | 1.968 | 1.88 | 1.934 | 2.019 |
| **Center, Layer 3** | 1.956 | 1.88 | 2.031 | 1.758 |

Figure S8: Sparsity of the mappings $\Pi$ calculated in pairwise and center alignment by PASTE. We report the average number of nonzero values per row.

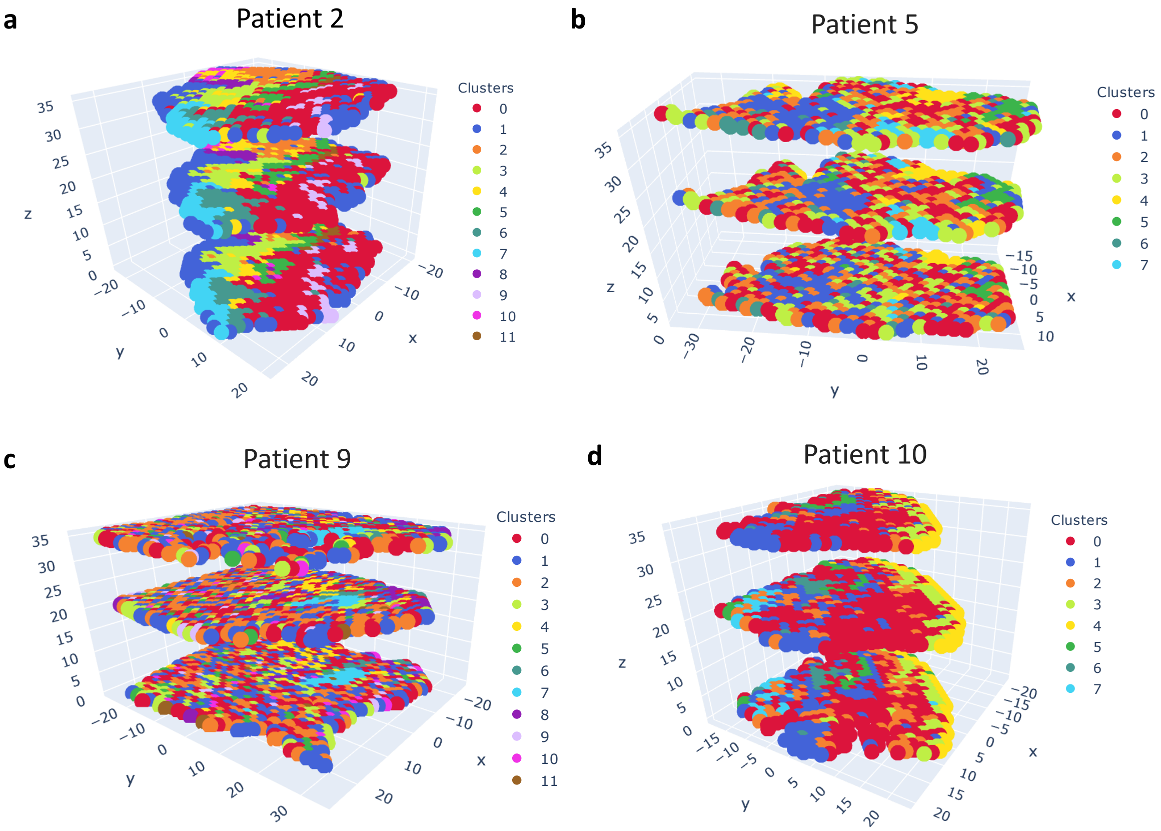


Figure S10: 3D layers alignment results on SCC data. Layers are color coded according to published cluster labels from [2]. The spatial coordinates of layers were aligned using mappings calculated by PASTE with Procrustes analysis (Section S1.2). The the $x, y$ coordinates are in 0.1mm scale while the scale of $z$ coordinate was changed for illustrative purposes.

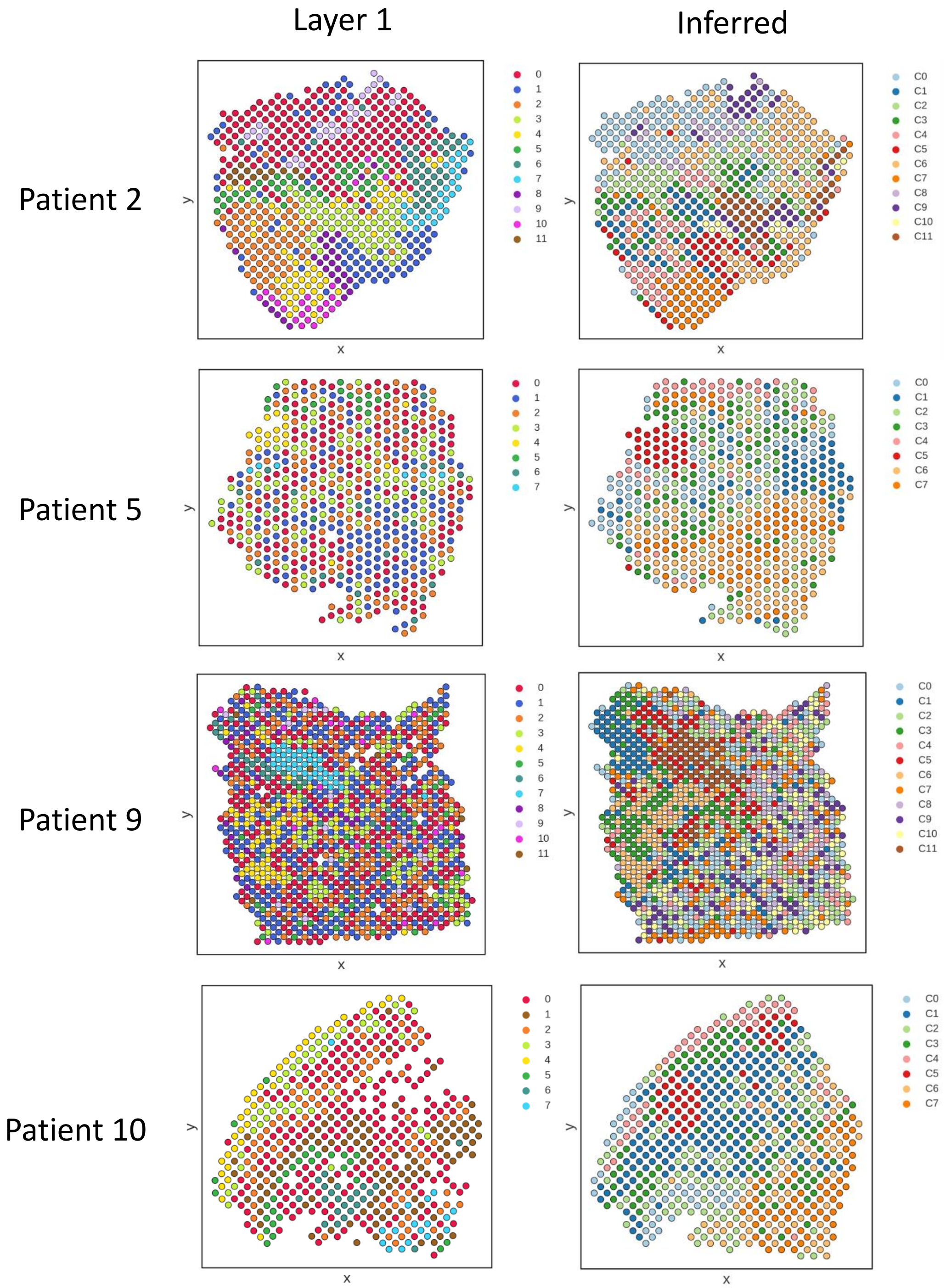


Figure S11: Comparison of published clusters and clusters obtained by PASTE on ST data from SCC patients 2, 5, 9, and 10 in [2]. (Left) The published cluster labels from [2] of spots in layer 1 from each of the four patients. (Right) $K$-means clustering of inferred center layer from PASTE.

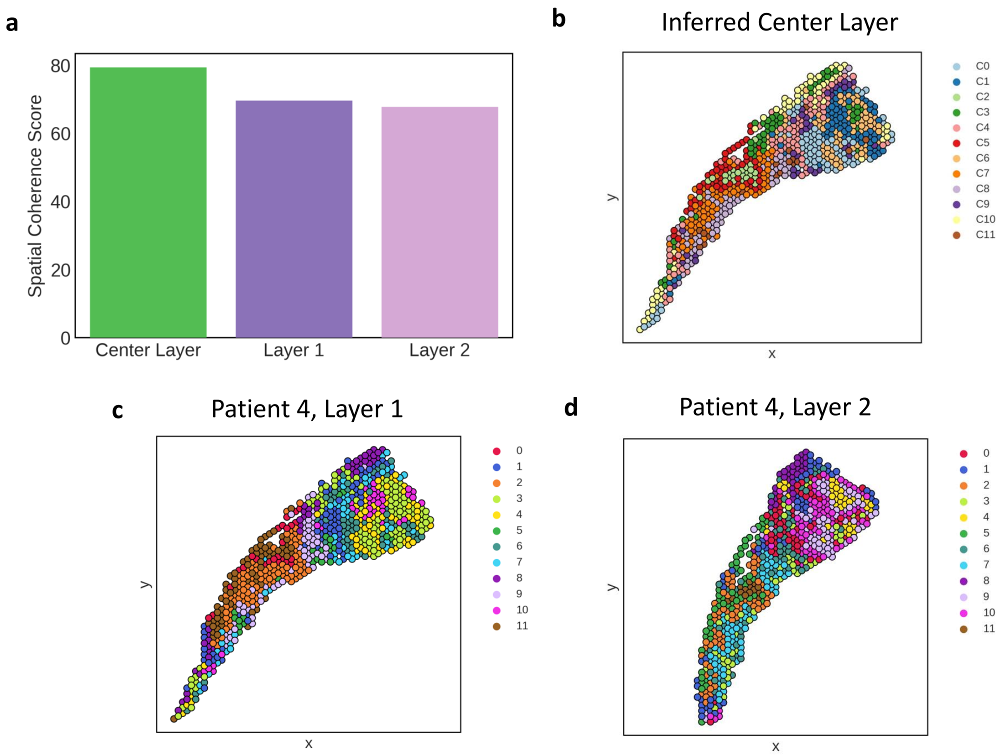


Figure S12: Center layer integration results on SCC Visium data [2]. (a) Spatial coherence comparison between center layer and real layers for patient 4. (b) Distribution of cluster-labeled spots in the inferred center layer of patient computed by PASTE. (c) Distribution of cluster-labeled spots in layer 1 of patient 4. (d) Distribution of cluster-labeled spots in layer 2 of patient 4.

### References

[1] Brian Hie, Bryan Bryson, and Bonnie Berger. Efficient integration of heterogeneous single-cell transcriptomes using scanorama. *Nature Biotechnology*, 37(6):685–691, 2019.

[2] Andrew Ji, Adam Rubin, Kim Thrane, Sizun Jiang, David Reynolds, Robin Meyers, Margaret Guo, Benson George, Annelie Mollbrink, Joseph Bergenstråhle, Ludvig Larsson, Yunhao Bai, Bokai Zhu, Aparna Bhaduri, Jordan Meyers, Xavier Rovira-Clavé, S Hollmig, Sumaira Aasi, Garry Nolan, and Paul Khavari. Multimodal analysis of composition and spatial architecture in human squamous cell carcinoma. *Cell*, 182:1661–1662, 09 2020.

[3] W. Kabsch. A solution for the best rotation to relate two sets of vectors. *Acta Crystallographica Section A*, 32(5):922–923, Sep 1976.

[4] F. Markley and D. Mortari. Quaternion attitude estimation using vector observations. *Journal of The Astronautical Sciences*, 48:359–380, 2000.

[5] Patrik L Ståhl, Fredrik Salmén, Sanja Vickovic, Anna Lundmark, José Fernández Navarro, Jens Magnusson, Stefania Giacomello, Michaela Asp, Jakub O Westholm, Mikael Huss, et al. Visualization and analysis of gene expression in tissue sections by spatial transcriptomics. *Science*, 353(6294):78–82, 2016.

[6] Grace Wahba. A least squares estimate of satellite attitude. *SIAM Review*, 7(3):409–409, 1965.
